# Bidirectional modulation of pain-related behaviors in the zona incerta

**DOI:** 10.1101/2021.03.10.434888

**Authors:** Sudhuman Singh, Spring Valdivia, Omar Soler-Cedeño, Anisha P. Adke, Barbara Benowitz, Daniela Velasquez, Torri D. Wilson, Yarimar Carrasquillo

## Abstract

Central amygdala neurons expressing protein kinase C-delta (CeA-PKCδ) are sensitized following nerve injury and promote pain-related responses in mice. The neural circuits underlying modulation of pain-related behaviors by CeA-PKCδ neurons, however, remain unknown. In this study, we identified a functional monosynaptic inhibitory neural circuit that originates in CeA-PKCδ neurons and terminates in the ventral region of the zona incerta (ZI), a subthalamic structure previously linked to pain processing. Behavioral experiments further show that chemogenetic inhibition of GABAergic ZI neurons is sufficient to induce bilateral hypersensitivity in uninjured mice as well as contralateral hypersensitivity after nerve injury. In contrast, chemogenetic activation of GABAergic ZI neurons reverses nerve injury-induced hypersensitivity, demonstrating that silencing of the ZI is required for injury-induced behavioral hypersensitivity. Our results identify a previously unrecognized inhibitory efferent pathway from CeA-PKCδ neurons to the ZI and demonstrate that ZI-GABAergic neurons can bidirectionally modulate pain-related behaviors in mice.

## Introduction

Persistent pain resulting from lesions or diseases affecting the peripheral and central nervous system can severely affect a person’s life over time if left untreated (Dworkin, 2002; Treede et al., 2008). Understanding the neural circuits underlying pain processing and how they are recruited in a maladaptive manner following injury is crucial for the development of improved treatment options for persistent pain. Several neuroimaging, pharmacological and electrophysiological studies in humans and animals demonstrate that the amygdala is a key locus in persistent pain processing (Bernard and Besson, 1990; Zald, 2003; Neugebauer et. al., 2004; Carrasquillo and Gereau, 2007; Bushnell et. al., 2013). A recent study further demonstrated that the CeA can both enhance and decrease pain-related behaviors in a cell-type-specific manner (Wilson et al., 2019). CeA neurons expressing protein kinase C-delta (CeA-PKCδ), for example, are sensitized by nerve injury and promote pain-related responses. In contrast, neurons expressing somatostatin are inhibited by nerve injury and promote decreases in pain-related behaviors. The circuit and cellular mechanisms responsible for bidirectional modulation of pain-related responses in the CeA, however, are still unclear.

In the present study, we began to address this question by characterizing the efferent projections from CeA-PKCδ neurons. Our cell-type-specific anatomical experiments identified the zona incerta (ZI) as one of the efferent targets of CeA-PKCδ neurons. The ZI is a subthalamic nucleus located ventrolateral to the medial lemniscus and dorsomedial to the substantia nigra (Ricardo,1981). The ZI is comprised of heterogeneous groups of cells defined by the expression of molecular markers such as parvalbumin, tyrosine hydroxylase, somatostatin, calbindin, and glutamate (Mitrofanis, 2005). The functions of the ZI are not fully understood but recent studies suggest that this brain region contributes to fear conditioning (Zhou et al., 2018) binge eating (Zhang and Van Den Pol, 2017), defensive behaviors (Chou et al., 2018) and predatory hunting (Zhao et al., 2019). In rodent models of pain, changes in neural activity have been reported in the ZI (Masri er al., 2009) and behavioral studies further show that experimentally modulating the activity of ZI neurons alters behavioral hypersensitivity (Petronilho et. al., 2012; Moon and Park, 2017; Hu et. al., 2019; Wang et. al., 2020). Most of the literature suggests, for example, that the ZI is inhibited in the context of pain and that inhibition drives behavioral hypersensitivity (Masri et. al., 2009; Moon et. al., 2016; Moon and Park, 2017; Hu et. al., 2019). A recent study, however, suggests the opposite- the ZI is activated in the context of pain and increases in neuronal activity drive hypersensitivity (Wang et. al., 2020). These seemingly conflicting results suggest that modulation of pain in the ZI is complex and, most likely, cell-type and circuit-specific. Identifying the sources of excitation and inhibition of ZI neurons in the context of pain will be important to begin untangling the mechanisms underlying modulation of pain in the ZI.

Based on our anatomical findings demonstrating a projection from CeA-PKCδ neurons to the ZI, in combination with previous work showing that CeA-PKCδ neurons are GABAergic and display increases in activity following injury, we hypothesized that inhibitory inputs from CeA-PKCδ neurons are a source of pain-related inhibition in the ZI that results in increases in pain-related behaviors. In the present study, we tested this hypothesis using a combination of cell-type-specific anatomical traces, opto-assisted circuit mapping, chemogenetic manipulations and behavioral assays to measure hypersensitivity in mice. Our combined results show that there is a functional monosynaptic inhibitory efferent pathway from CeA-PKCδ neurons to the ZI and further demonstrate that ZI-GABAergic neurons can bidirectionally modulate pain-related behaviors in mice.

## Results

### Identification of CeA-PKCδ neuronal efferent targets

The neural pathways underlying modulation of pain-related behaviors by CeA-PKCδ neurons remains unknown. To begin to address this question, we stereotaxically injected a cre-dependent adeno-associated virus (AAV) anterograde tracer (ChrimsonR or mCherry) into the CeA of *Prkcd*-cre mice (**Figure 1A**). The transduction of ChrimsonR or mCherry in CeA-PKCδ cells was confirmed with immunostaining for PKCδ (**Figure 1B**). Mapping of ChrimsonR (or mCherry) positive axonal terminals revealed CeA-PKCδ efferent projections with dense, moderate and sparse labeling in 17 brain regions throughout the brain, including the basal forebrain, striatum, thalamus, hypothalamus, midbrain, pons and medulla (**Figure 1C-D** and **Table 1**). All 17 brain regions identified in our experiments have been previously defined as output regions of the CeA using traditional anterograde tracers (Shinonaga et al., 1992; Reardon and Mitrofanis, 2000; Barbier et al., 2017; Zhou et al., 2018; Aggleton, 2000), validating our experimental approach to study the efferent projections of CeA-PKCδ neurons.

**Table 1.** CeA-PKCδ Neuronal Efferent Targets. Semi-quantitative analysis of the density of axonal terminals in brain regions from 5 PKCδ-Cre mice stereotaxically injected with a cre-dependent adeno-associated virus anterograde tracer (ChrimsonR, mCherry, EGFP) into the CeA. Rightmost column is from experiment 265945645 of the Mouse Brain Connectivity Atlas of the Allen Brain Institute (http://connectivity.brain-map.org/). - no expression; + sparce; ++ moderate; +++ dense

| Area | Abbreviations | ET 987<br>(ChrimsonR) | ET 903<br>(ChrimsonR) | ET832<br>(mCherry) | ET835<br>(mCherry) | Allen Brain Atlas<br>(EGFP) |
| --- | --- | --- | --- | --- | --- | --- |
| **Striatum and Basal Forebrain** |  |  |  |  |  |  |
| Bed nucleus of stria terminalis | BNST | +++ | +++ | +++ | +++ | +++ |
| Globus pallidus | GP | + | + | ++ | + | + |
| Extended amygdala | EA | +++ | +++ | +++ | + / ++ | +++ |
| Central amygdala | CeA | +++ | +++ | ++ | + / ++ | ++ |
| Substantia innominata | SI | ++ | ++ | ++ | + / ++ | ++ |
| **Thalamus** |  |  |  |  |  |  |
| Subthalamic nucleus | STh | ++ | + | + | - | - |
| Zona Incerta | ZI | ++ | + | ++ | - | + |
| Para subthalamic nucleus | PSTh | ++ | ++ | + / ++ | + | - |
| **Hypothalamus** |  |  |  |  |  |  |
| lateral preoptic area | LPO | + | + | + / ++ | + | + |
| Lateral hypothalamus | LH | + | + | + / ++ | + | + |
| **Midbrain** |  |  |  |  |  |  |
| Ventral tegmental area | VTA | + / ++ | + | + / ++ | - | - |
| Substantia nigra | SN | ++ | + / ++ | ++ | + | - |
| Pedunculo pontine tegmental nucleus | PTg | + | + | ++ | - | - |
| Laterodorsal tegmental nucleus | LDTg | + | + | - | + | - |
| Periaqueductal grey | PAG | + / ++ | + | - | + | - |
| Reticular formation | mRT | + / ++ | + | + / ++ | + / ++ | - |
| **Pons** |  |  |  |  |  |  |
| Lateral parabrachial | LPB | +++ | +++ | ++ / +++ | + | - |
| Locus coeruleus | LC | + | + | ++ | + | - |

Semi-quantitative analysis, performed by visual examination of high magnification images, further revealed that axonal terminals in the output regions of CeA-PKCδ neurons have different terminal densities and organization patterns. Most of the brain regions identified had either few or moderate numbers of terminals, with only three regions, including the bed nucleus of stria terminalis, extended amygdala and parabrachial nucleus, containing high densities of labeling (**Figure 1C-D** and **Table 1**).

As summarized in **Table 1**, dense to moderate labeling was consistently seen in the bed nucleus of stria terminalis, extended amygdala and central amygdala of all 5 brains analyzed. Dense labeling was also seen in the lateral parabrachial nucleus of 3 of the 5 brains analyzed, with sparse labeling observed in 1 brain and no labeling in the other brain. Moderate to sparse labeling was consistently observed in the substantia innominata in all 5 brains; in the zona incerta, para-subthalamic nucleus, substantia nigra and reticular formation in 4 of 5 brains; and in the subthalamic nucleus and ventral tegmental area in 3 of the 5 brains analyzed. Lastly, sparce labeling was consistently seen in the globus pallidus, lateral preoptic area and lateral hypothalamus in all 5 brains; in the locus coeruleus in 4 of 5 brains and in the pedunculopontine tegmental nucleus, laterodorsal tegmental nucleus and periaqueductal grey in 3 of the 5 brains. Consistent with previous studies using traditional anterograde tracers in the CeA (Shinonaga et al., 1992; Reardon and Mitrofanis, 2000; Barbier et al., 2017; Zhou et al., 2018; Aggleton, 2000), no terminal labeling was observed in cortical regions of any of the 5 brains evaluated.

Mapping of the injection sites in all 5 brains shows that all injections were restricted to the CeA (**Figure 1 – figure supplement 1**). The number of transduced neurons in these mice were comparable to the expression levels of PKCδ-tdTomato neurons in the CeA of *Prkcd*-cre::Ai9 mice, demonstrating robust transduction efficiency in these experiments. Differences in the size and rostrocaudal distributions of the injections, however, might explain the differences observed in efferent projections between brains.

**Figure 1.**
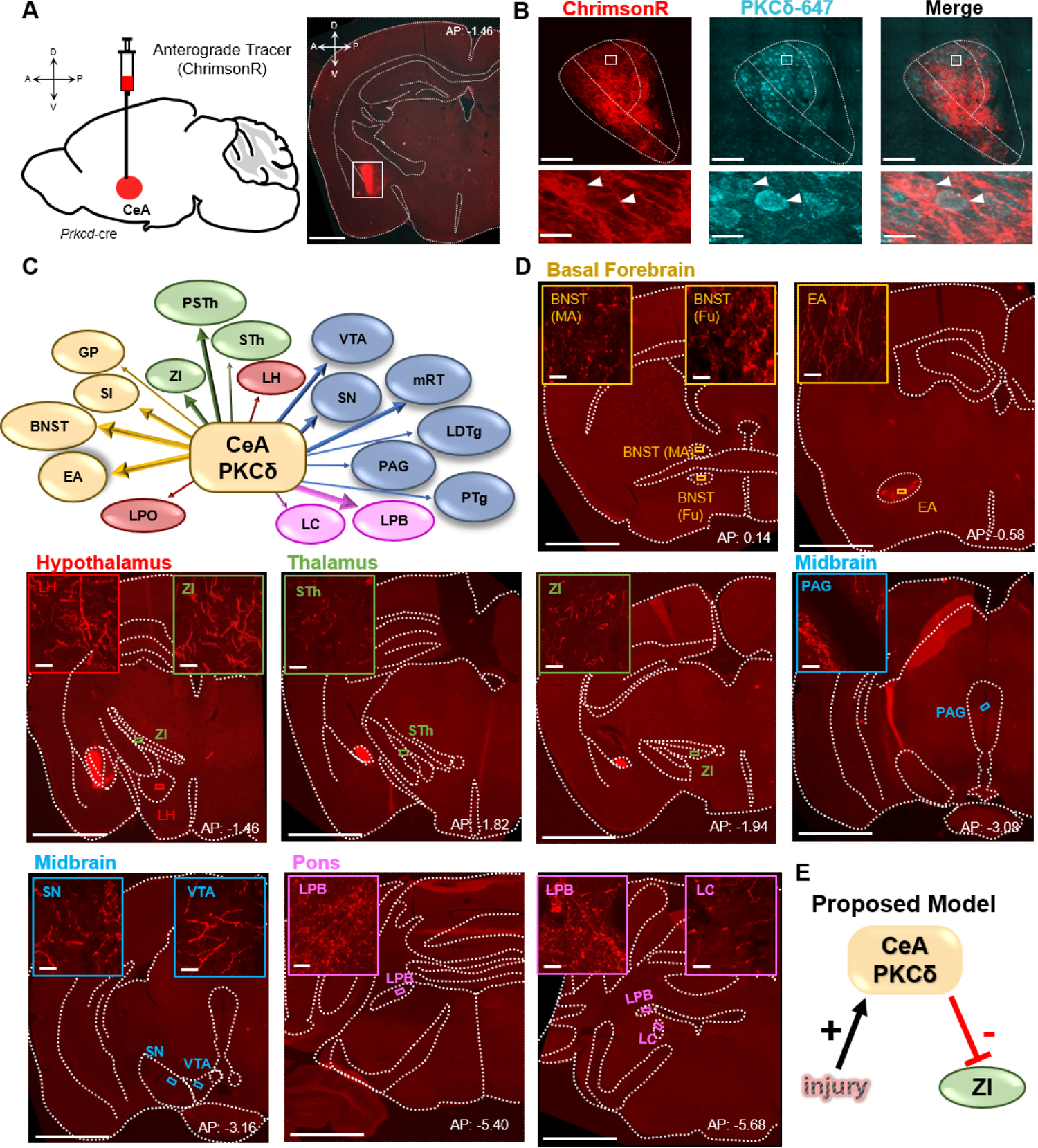
CeA-PKCδ neurons efferent targets. **(A)** Schematic of experimental approach. ChrimsonR-tdTomato was unilaterally injected into the CeA of a PKCδ-cre mouse. A representative coronal brain slice of an injected mouse is shown on the right panel, with ChrimsonR-tdTomato shown in red. Scale bar represents 1000 µM. **(B)** Representative high magnification images of the CeA in a coronal brain slice of a ChrimsonR-tdTomato injected mouse. ChrimsonR-transduced cells are shown in red and neurons immunostained for PKCδ in cyan. The merged image is shown on the right panel. Lower insets depict higher magnification images of the areas delineated by the white box in the upper images. White arrowheads highlight representative transduced cells that are also positive for PKCδ. Scale bars represent 100 µM for low magnification and 10 µM for high magnification images. **(C)** Summary diagram illustrating CeA-PKCδ neuron efferent projections within the brain. Forebrain regions are shown in yellow, hypothalamic structures in red, thalamus in green, midbrain in blue and pons in fuchsia. The thickness of the arrows depict the density of labeling (sparse, moderate or dense). (**D**) Low magnification representative images of brain regions with axonal terminals from CeA-PKCδ cells. Insets in each image are high magnification images depicting axonal terminals within the regions delineated by the boxes in the respective low magnification images. Scales are 1000 µM for low magnification images and 20 µM for high magnification images. bed nucleus of stria terminalis medial (BNST-MA); bed nucleus of stria terminalis fusiform nucleus (BNST-Fu); extended amygdala (EA); substantia innominate (SI); lateral preoptic area (LPO); globus pallidus (GP); lateral hypothalamus (LH); Subthalamic nucleus (STh); Zona incerta (ZI); Parasubthalamic nucleus (PSTh); periaqueductal grey (PAG); substantia nigra (SNR); ventral tegmental area (VTA); pedunculopontine tegmental nucleus (PTg); laterodorsal tegmental nucleus (LDTg); reticular formation (mRT); lateral parabrachial (LPB); locus coeruleus (LC). **See Figure1 – figure supplement 1.**

### CeA-PKCδ neurons send monosynaptic inhibitory projections to the Zona Incerta

The ZI was among the regions identified as an efferent target of CeA-PKCδ neurons in our anatomical experiments (**Figure 1C**). This subthalamic brain region was of interest because previous studies have shown that reduced activity in GABAergic neurons of the ZI correlates with pain-related behaviors (Masri, et al., 2009; Moon et al., 2016; Moon and Park, 2017; Hu et al., 2019). Given that CeA-PKCδ neurons are GABAergic and are activated in the context of pain (Wilson et al., 2019), we hypothesized that pain-related inhibition of the ZI is mediated by injury-induced activation of CeA-PKCδ neurons (**Figure 1E**). We began to test this hypothesis by first validating the projections from CeA-PKCδ neurons to the ZI using a retrograde tracer approach where the retrograde tracer cholera toxin B (CTB), conjugated with Alexa Fluor 647, was stereotaxically injected into the ZI of *Prkcd*-cre::Ai9 mice (**Figure 2A**). As illustrated in **Figure 2B**, the retrograde tracer CTB-647 was readily detected in CeA-PKCδ neurons after injection into the ZI, validating the existence of an anatomical pathway from CeA-PKCδ neurons to the ZI.

**Figure 2.**
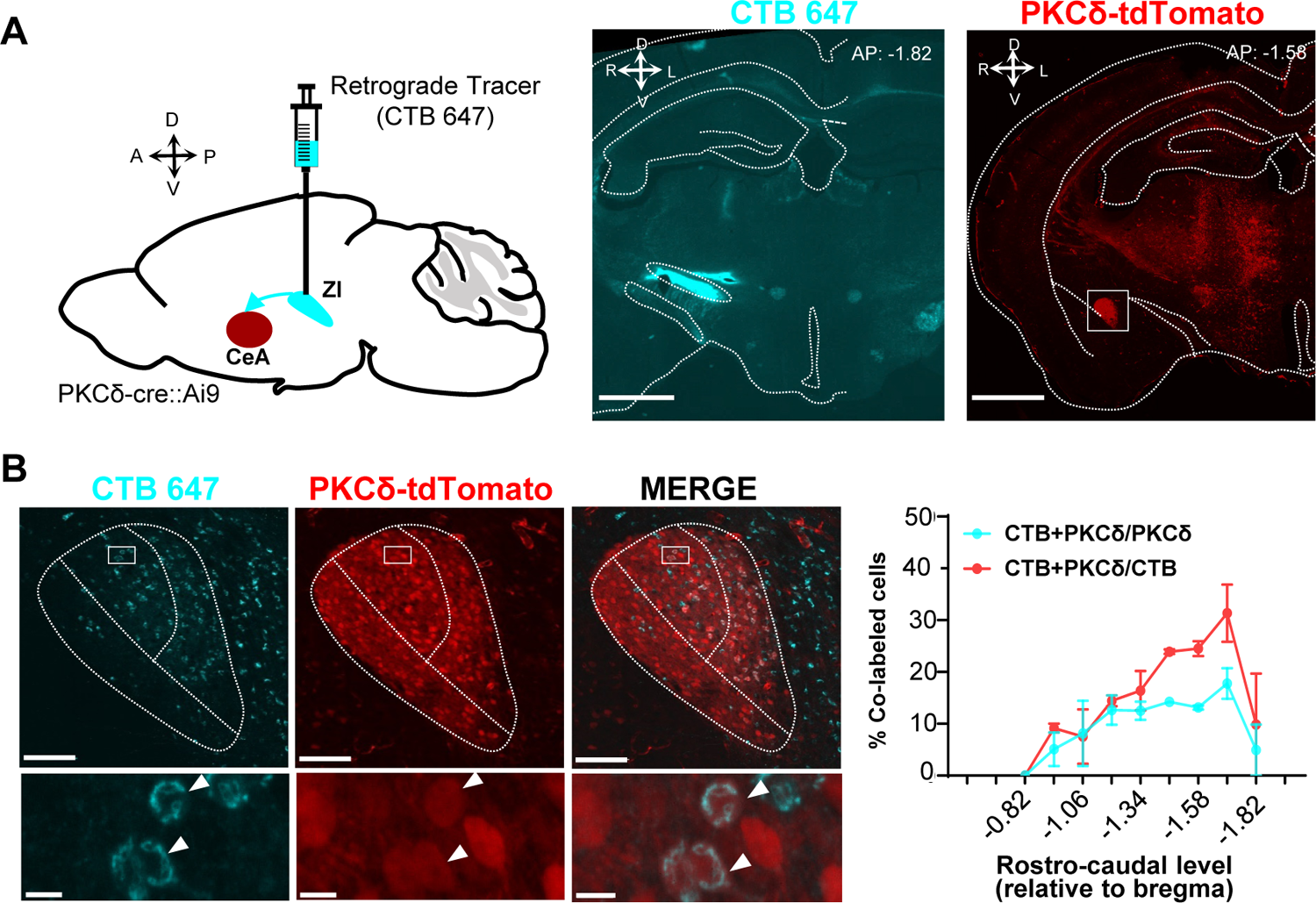
Retrograde tracing of CeA-PKCδ projections to the ZI. (**A**) Schematic drawing of experimental approach (left panel). Cholera toxin B (CTB-647) was injected into the ZI of a PKCδ-cre::Ai9 mouse brain. A representative coronal brain slice depicting the focal injection of CTB-647 (cyan) into the ZI is shown in the middle panel. A representative coronal brain slice containing the CeA is shown in the right panel. PKCδ-tdTomato cells are shown in red. The white square delineates the area magnified in panel B. Scale bar represents 1000 µM. (**B**) Representative high magnification images of the CeA in a PKCδ-cre::Ai9 mouse injected with CTB-647 in the ZI. CTB-positive cells are shown in cyan and PKCδ-tdTomato cells in red. The merged image is shown on the right. Lower insets show higher magnification images of the area delineated by the white squares in the top panel. Arrowheads highlight cells that are positive for CTB and PKCδ-tdTomato. Scale bars represent 100 µM (top panel) and 10 µM (bottom panel). The mean ± SEM percentage of cells co-labeled for PKCδ and CTB as a function of the rostro caudal level is shown on the right (n=2 mice, 8 slices per mouse).

To characterize the functional connectivity between CeA-PKCδ neurons and ZI GABAergic neurons, we stereotaxically injected AAV-hsyn-ChR2-EYFP into the CeA of VGAT-cre::Ai9 mice and performed patch-clamp recordings in acute brain slices containing the ZI (**Figure 3A**). The stable expression of ChR2-EYFP in the CeA is indicated by the presence of green fluorescent signal in the peri-somatic region of VGAT-positive neurons in CeA. Consistent with our anatomical findings, tracing of the EYFP-labeled axonal terminals revealed moderate labeling in both VGAT-positive and VGAT-negative ZI neurons. Optogenetic stimulation of ChR2-expressing CeA terminals in the ZI with blue light further showed robust inhibitory post-synaptic currents in 60% VGAT-positive (12 out of 20 cells) and 53% VGAT-negative (8 out of 15 cells) ZI neurons (**Figure 3B**). These optically evoked postsynaptic responses occurred in the presence of TTX and 4-AP, demonstrating that the inputs from the CeA to the ZI are monosynaptic.

**Figure 3.**
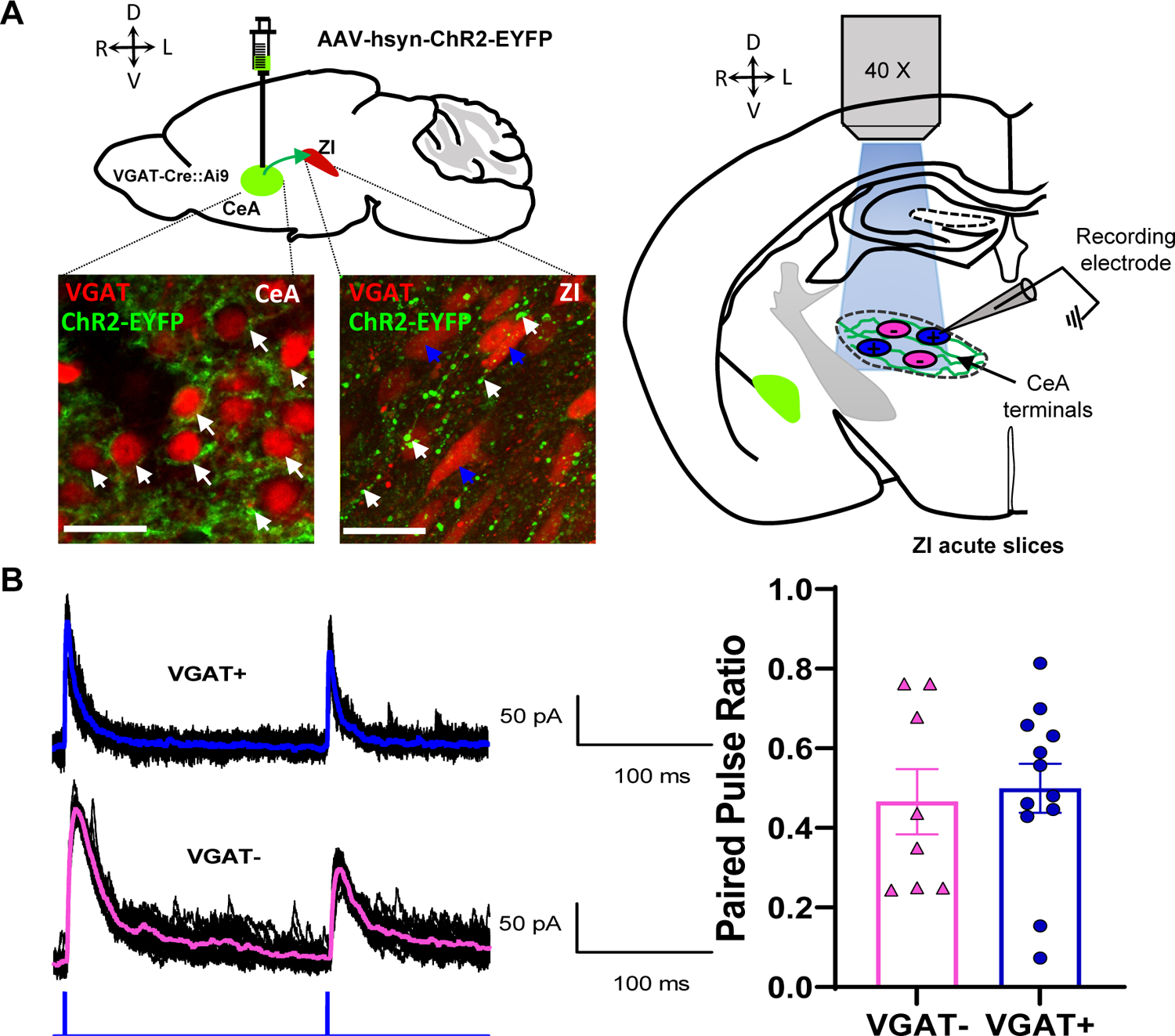
Optogenetic activation CeA terminals in the ZI results in robust inhibitory postsynaptic responses. (**A**) Schematics of the experimental approach. VGAT-cre::Ai9 mice were stereotaxically injected with AAV-hsyn-hChR2-EYFP into the CeA. After 4 weeks, ChR2-EYFP is expressed in CeA neurons (perisomatic, white arrows, lower left panel) and in the ZI (CeA terminals, white arrows, lower right panel) in proximity to VGAT positive (blue arrows) cells. Scale bars are 20 µM. Schematic diagram for ex-vivo whole-cell recordings in acute ZI brain slices is shown in the right panel. Responses of VGAT-positive (blue) and VGAT-negative (magenta) cells are recorded upon optically activating ChR2-expressing CeA terminals. (**B**) Representative traces showing evoked inhibitory postsynaptic currents in ZI VGAT-positive and VGAT-negative neurons in response to paired pulse stimulation of CeA terminals (5 ms duration, 200 ms inter stimulus interval). The mean +/- SEM paired pulse ratio is shown on the right panel (n=8 VGAT-negative and 12 VGAT-positive cells).

### Inhibition of ZI-GABAergic neurons is sufficient to induce bilateral hypersensitivity

Previous studies have shown that ZI-GABAergic neurons are inhibited in the context of pain (Masri et al., 2009; Moon et al., 2016; Moon and Park 2017; Hu et al., 2019). To establish a causal link between reduced activity of ZI-GABAergic neurons and pain-related behaviors, we utilized a chemogenetic approach coupled with a battery of pain behavioral assays to measure tactile and thermal sensitivity in naïve mice as well as in mice with the sciatic nerve cuff model of neuropathic pain. Whole-cell current-clamp recordings in acute ZI slices prepared from VGAT-cre mice stereotaxically injected with a cre-dependent AAV encoding the inhibitory designer receptors exclusively activated by designer drugs (G_i_ DREADD) hM4Di into the ZI was used to validate hM4Di-mediated inhibition of ZI neurons (**Figure 4A**). As illustrated in **Figure 4A**, bath application of CNO (10 µM) significantly inhibited firing responses in hM4Di-transduced neurons, with no measurable effect observed in response to bath application of the saline vehicle control. Histological verification of injection sites at the end of the experiments further demonstrated that transduction of hM4Di-mCherry was restricted to the ZI (**Figure 4B** and **Figure 4 – figure supplement 1**). The numbers and rostrocaudal distribution of transduced cells within the ZI of mice stereotaxically injected with hM4Di-mCherry was comparable to the numbers and rostrocaudal distribution of VGAT-positive cells in the ZI of VGAT-Cre::Ai9 mice, demonstrating robust transduction efficiency (**Figure 4C**).

**Figure 4.**
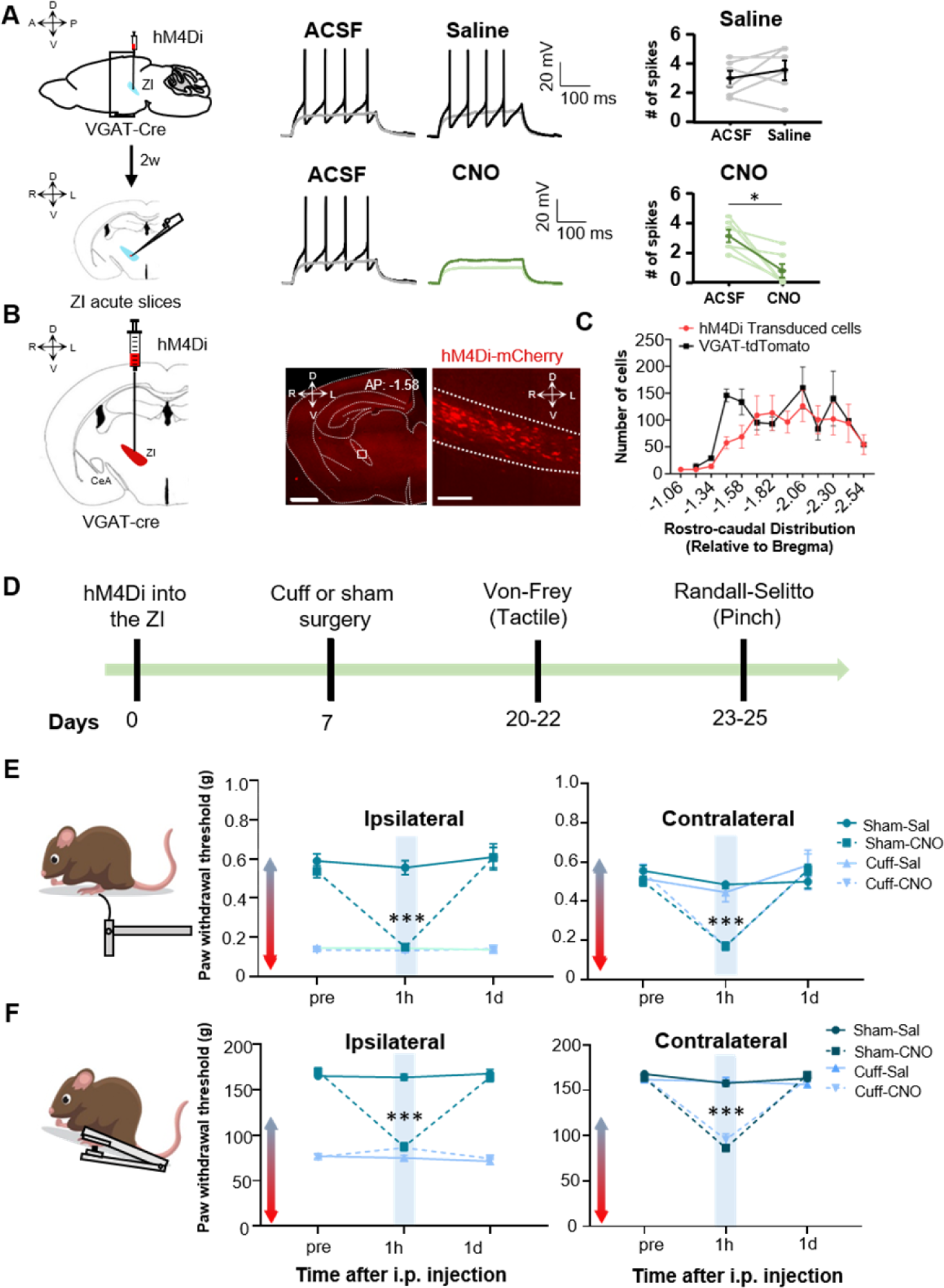
Inhibition of ZI GABAergic neurons is sufficient to induce bilateral tactile and pressure hypersensitivity in uninjured mice. (**A**) Schematic of the experimental approach. VGAT-Cre mice were stereotaxically injected with AAV-DIO-hM4Di-mCherry into the ZI. Current-clamp recordings were obtained from mCherry-positive cells in acute ZI slices 2 weeks after the injection. Representative traces of whole-cell current-clamp recording obtained from ZI neurons transduced with hM4Di-mCherry before (left) and after (right) bath application of 10 µM CNO (lower panel) or vehicle (top panel). Action potential were elicited using 500-ms depolarizing current injection that evoked 2 to 5 action potential before bath the application. The same amplitude of depolarizing current injection was used before and after bath application. Summary graphs depicting the mean ± SEM number of spikes before and after bath treatment are shown on the right panel (n = 6 neurons per treatment; *p<0.05 for ACSF vs CNO). (**B**) Schematic diagram for unilateral stereotaxic injection of AAV-DIO-hM4Di-mCherry in ZI region of VGAT-Cre mice. A representative image of a coronal mouse brain slice from a VGAT-Cre mouse injected with AAV-DIO-hM4Di-mCherry into the ZI is shown on the middle panel. The area delineated by the white rectangle in the middle panel is shown at higher magnification in the right panel, with mCherry positive cells shown in red. Scale bars represent 1000 µm (left) and 100 µm (right). (**C**) Mean ± SEM number of hM4Di-transduced cells and VGAT-tdTomato labeled cells in the ZI as a function of rostrocaudal level relative to bregma (n=11 mice for hM4Di-transduced neurons; n=4 mice for VGAT-tdTomato neurons). (**D**) Timeline for behavioral experiments. (**E-F**) Von Frey (**E**) and Randall Selitto (**F**) responses shown as mean ± SEM paw withdrawal threshold in the ipsilateral (left panel) and contralateral (right panel) hindpaw before, 1 h and 1 day after CNO or vehicle i.p. injection in cuff or sham mice stereotaxically injected with AAV-DIO-hM4Di-mCherry into the ZI. (n=8 mice per treatment; ***p<0.0001 for pre-injections vs 1 h after CNO in sham-hM4Di CNO treatment for both ipsi and contralateral hindpaws and for cuff-hM4Di-CNO treatment in the contralateral hindpaw; two-way ANOVA followed by Dunnett’s multiple comparisons test). See **Figure 4 – figure supplement 1**.

The effects of selective chemogenetic inhibition of VGAT-positive ZI cells on pain-related responses to tactile and pressure stimulation of the hindpaws were measured before and after i.p. injection of CNO to activate the G_i_DREADDs, or saline as the vehicle control in both cuff and sham mice (**Figure 4D**). As expected, following cuff implantation on the sciatic nerve, tactile and pressure sensitivity was significantly lower in the hindpaw ipsilateral to cuff implantation compared to the contralateral hindpaw or both hindpaws in sham treated mice (**Figures 4E-F**). As illustrated in **Figures 4E-F**, chemogenetic inhibition of VGAT-positive ZI neurons resulted in robust bilateral hypersensitivity to tactile and pressure stimulation in sham mice as well as contralateral hypersensitivity in cuff-implanted mice. Thus, compared to pre-injection values, paw withdrawal thresholds in response to tactile or pinch stimulation were significantly (p < 0.0001) reduced bilaterally following CNO injections in sham mice. Similarly, withdrawal thresholds in the hindpaw contralateral to cuff implantation were significantly (p < 0.0001) reduced after CNO injection, compared to pre-injection thresholds.

Withdrawal thresholds in the hindpaw ipsilateral to cuff treatment were indistinguishable before and after selective chemogenetic inactivation of VGAT-positive ZI neurons with CNO in hM4Di ZI-injected mice, demonstrating that inhibition of ZI cells does not measurably affect cuff-induced hypersensitivity to tactile and pressure stimulation. Notably, paw withdrawal thresholds in sham mice and the contralateral hindpaw of cuff-implanted mice following the chemogenetic inhibition of ZI VGAT-positive neurons were comparable to the withdrawal thresholds measured in the ipsilateral hindpaw of cuff-implanted mice, demonstrating that inhibition of these neurons is sufficient to elicit hypersensitivity in the absence of injury that resembles the hypersensitivity observed following sciatic nerve cuff implantation. The effects of chemogenetic inhibition of VGAT-positive ZI neurons on tactile and pressure sensitivity were transient, as paw withdrawal thresholds in all treated animals returned to pre-injection values 1 day following the CNO injections. Importantly, tactile and pressure sensitivity was unaltered in saline-injected mice, demonstrating that the CNO-induced effects were not due to handling or hM4Di expression (**Figures 4E-F**).

Previous studies have demonstrated that modulation of pain-related behaviors in the CeA, including by CeA-PKCδ neurons, is modality-dependent (Wilson et., 2019). In order to evaluate if modulation of hypersensitivity in the ZI is also modality-specific, the next set of experiments assessed the effects of inhibition of ZI VGAT-positive neurons on heat and cold sensitivity in both sham and cuff-implanted mice using the Hargreaves and acetone evaporation tests, respectively. As illustrated in **Figure 5**, cuff implantation in the sciatic nerve resulted in hypersensitivity to both heat and cold stimulation in the hindpaw ipsilateral to cuff implantation compared to the contralateral hindpaw or either hindpaw in sham treated mice. Behavioral responses to cold and heat stimulation of the hindpaws, however, were unaltered by chemogenetic inhibition of VGAT-positive neurons in the ZI in all animals tested. Taken together, the results from these chemogenetic experiments demonstrate that inhibition of VGAT-positive neurons in the ZI is sufficient to induce hypersensitivity in the absence of injury in a modality-specific manner.

**Figure 5.**
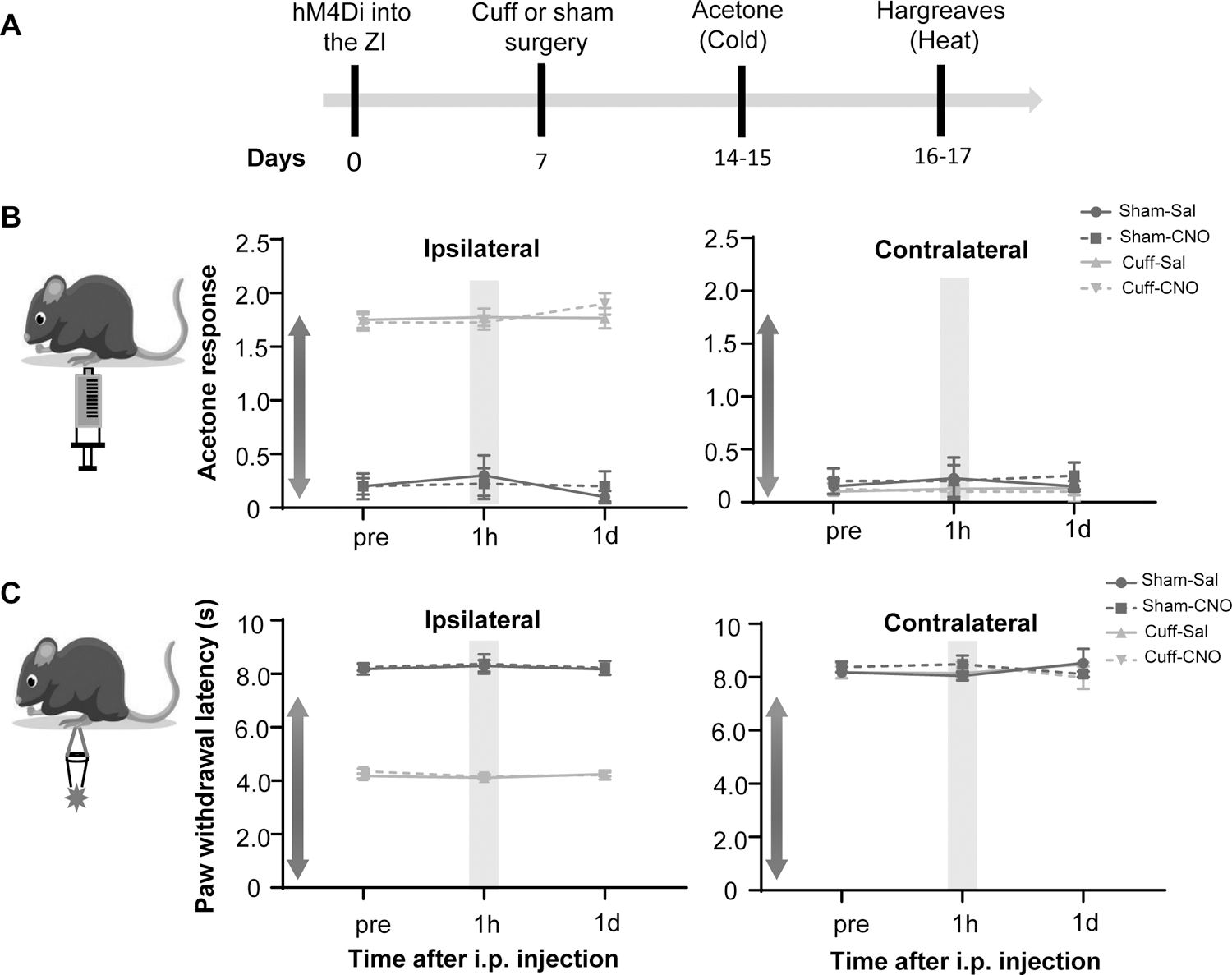
Cuff-induced thermal hypersensitivity is unaltered by chemogenetic inhibition of ZI-VGAT cells. (A) Timeline for behavioral experiments. (**B-C**) Mean ± SEM acetone response (**B**) or paw withdrawal latency obtained from Hargreaves test (**C**) in the ipsilateral (left panel) and contralateral (right panel) hindpaw before, 1 h and 1 day after CNO or vehicle i.p. injection in cuff or sham mice stereotaxically injected with AAV-DIO-hM4Di-mCherry into the ZI. (n=8 mice per treatment; no significant differences were observed in any treatment; two-way ANOVA followed by Dunnett’s multiple comparisons test).

### Activation of ZI-GABAergic neurons reverses cuff-induced hypersensitivity to pinch but not thermal stimulation

The next set of experiments aimed to determine whether activation of GABAergic ZI neurons is sufficient to reverse cuff-induced hypersensitivity. We used a chemogenetic approach coupled with behavioral assays to measure tactile and heat sensitivity in mice following the implantation of a sciatic nerve cuff to model neuropathic pain. Histological verification of the injection sites at the end of the experiments demonstrated that transduction of hM3Dq-mCherry and control-mCherry was restricted to the ZI (**Figure 6A** and **Figure 6 – figure supplement 1 and 2**). The numbers and rostrocaudal distribution of transduced cells within the ZI were comparable in mice injected with hm3Dq-mCherry and control-mCherry and show robust transduction efficiency that is localized to the ZI (**Figure 6B**). CNO-mediated activation of neurons in VGAT-Cre mice stereotaxically injected with AAV8-DIO-hM3Dq-mCherry into the ZI was validated with immunohistochemical monitoring of c-Fos, the product of an immediate early gene that is commonly used as a marker of neuronal activity (**Figure 6C**). As illustrated in **Figure 6D**, i.p. injection of CNO resulted in robust c-Fos expression in the ZI of VGAT-Cre animals injected with hM3Dq, compared to the c-Fos expression observed in the ZI of saline-injected control mice that also expressed hM3Dq or CNO-injected control VGAT-Cre mice stereotaxically injected with the control virus (AAV8-DIO-mCherry). Quantification of ZI cells co-expressing c-Fos and mCherry further confirms that c-Fos expression in hM3Dq-transduced cells is significantly (p < 0.05) higher in CNO-treated mice than it is in saline-treated mice or in mCherry-transduced neurons from CNO-treated mice (**Figure 6E**).

**Figure 6.**
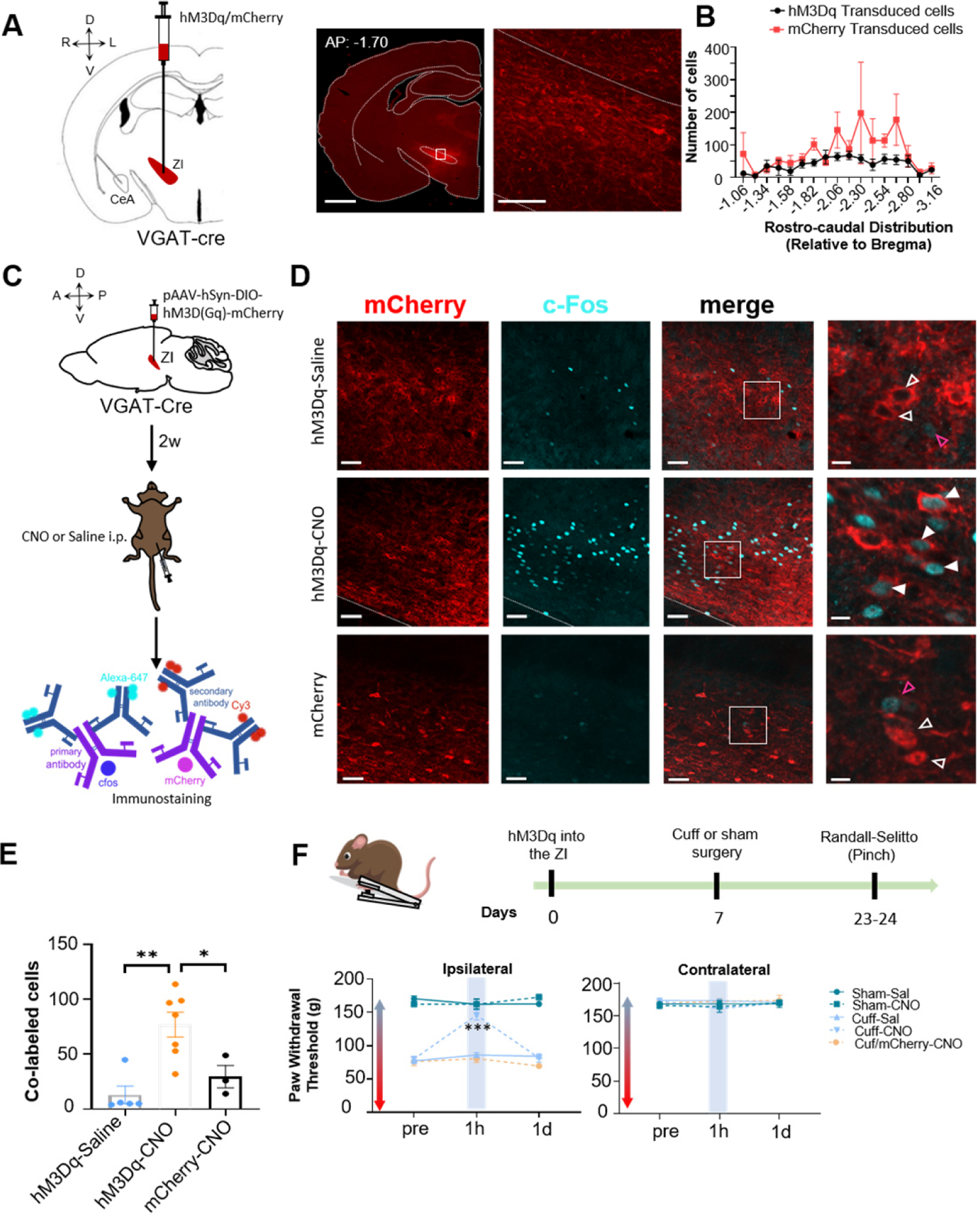
Activation of ZI GABAergic cells reverses cuff-induced hypersensitivity to pinch stimulation. **(A)** VGAT-Cre mice were injected with AAV-DIO-hM3Dq-mCherry or AAV-DIO-mCherry into the ZI. Low magnification representative image of a coronal brain slice shows the site of virus injection in red. The area delineated by the white rectangle is shown at higher magnification in the right image. Scale bars are 1000 µm for low magnification and 100 µm for high magnification images **(B)** Quantification of ZI cells transduced with hM3Dq and mCherry is shown as mean ± SEM (n=17 mice for hM3Dq transduced group and 7 mice for mCherry group). **(C)** c-Fos experimental timeline. **(D)** Representative images of coronal brain slices containing the ZI of VGAT-Cre mice injected with AAV-DIO-hM3Dq-mCherry (top and middle panels) or AAV-DIO-mCherry (bottom) into the ZI and i.p. treated with CNO (middle and bottom panels) or saline (top panel). Top panel shows images from mouse with mCherry (Control virus) injected in ZI. mCherry expression is shown in red and immunostaining for c-Fos in cyan. The merged images are shown in the rightmost panels. White boxes delineate the areas magnified on the right panel. Magenta open arrowheads point to cells that are positive for c-Fos only; white open arrowheads point to cells that are positive for mCherry only; solid arrowheads point to cells that are positive for both mCherry and c-Fos. Scale bars are 50 µm (low magnification) and 10 µm (high magnification). **(E)** Mean ± SEM numbers of c-Fos and mCherry transduced co-labeled cells per condition. (n=3-7 mice per condition; **p<0.01, *p<0.05; One-way ANOVA followed by Tukey’s multiple comparisons test). **(F)** Randall Selitto responses are shown as mean ± SEM paw withdrawal threshold in the ipsilateral (left panel) and contralateral (right panel) hindpaw before, 1 h and 1 day after CNO or vehicle i.p. injections in cuff or sham mice stereotaxically injected with AAV-DIO-hM3Dq-mCherry or AAV-DIO-mCherry into the ZI (n=6-8 mice per condition; ***p<0.0001 for pre-injections vs 1 h after CNO in cuff-hM3Dq CNO treatment; two-way ANOVA followed by Dunnett’s multiple comparisons test). **See Figure 6 – supplemental figures 1** and 2.

The effect of chemogenetic activation of VGAT-positive ZI cells on behavioral responses to pressure (pinching) stimulation of the hindpaws was measured before and after i.p. injection of CNO or saline in cuff and sham animals (**Figure 6F**). CNO-mediated activation of VGAT-positive ZI neurons led to significant (p < 0.0001) reversal of cuff-induced hypersensitivity in the hindpaw ipsilateral to cuff implantation, while no measurable effects were seen in saline-injected or CNO-injected mCherry-control mice. The effects of activation of VGAT-positive ZI neurons were specific to nerve injury as withdrawal thresholds in sham treated mice or the hindpaw contralateral to sciatic nerve cuff were comparable between groups. As illustrated in **Figure 7**, reversal of cuff-induced hypersensitivity was also modality-specific as behavioral responses to cold and heat stimulation of the hindpaws were unaffected by chemogenetic activation of VGAT-positive neurons in the ZI. Taken together, these results demonstrate that activation of VGAT-positive neurons in the ZI reverses cuff-induced pain hypersensitivity in a modality-specific manner.

**Figure 7.**
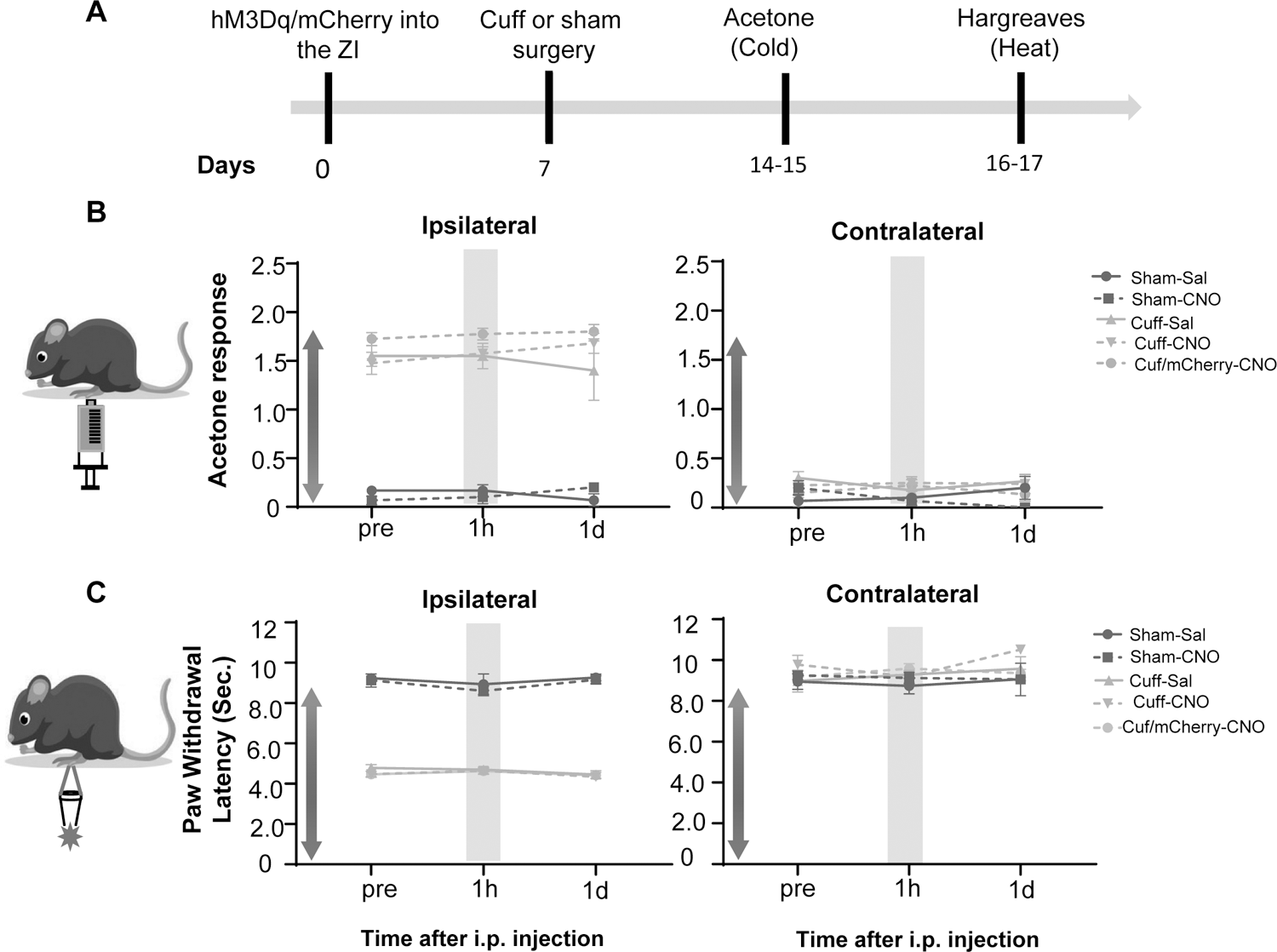
Cuff-induced thermal hypersensitivity is unaltered by chemogenetic activation of ZI-VGAT cells. **(A)** Experimental timeline. (**B-C**) Mean ± SEM acetone response (**B**) or paw withdrawal latency obtained from Hargreaves test (**C**) in the ipsilateral (left panel) and contralateral (right panel) hindpaw before, 1 h and 1 day after CNO or vehicle i.p. injection in cuff or sham mice stereotaxically injected with AAV-DIO-hM3Dq-mCherry or AAV-DIO-mCherry into the ZI. (n=4-8 mice per treatment; no significant differences were observed in any treatment; two-way ANOVA followed by Dunnett’s multiple comparisons test).

## Discussion

Signals for persistent neuropathic pain in distant parts of the body are perceived by the brain through ascending pathways, whereas the resulting responses to pain are heavily influenced by descending pathways from the brain. Maladaptive changes at any site in the pain neuraxis can result in persistent changes in pain perception. Nerve injury-induced increases in the activity of CeA-PKCδ neurons, for example, have been shown to contribute to behavioral hypersensitivity (Wilson et al., 2019). The neural circuits involved in modulation of nociceptive behaviors by CeA-PKCδ cells, however, remains unknown. In the present study, we identified 17 long-range efferent projections from CeA-PKCδ neurons, including the ZI which has been previously reported to be involved in pain processing. We further demonstrated that the ZI receives strong monosynaptic inhibitory inputs from CeA-PKCδ neurons and that chemogenetic manipulation of the activity of

GABAergic neurons in the ZI bidirectionally modulates pain-related behavioral outputs in a modality-specific manner. Altogether, our results identify the ZI as a functional anatomical target of the CeA that contributes to the modulation of pain-related behaviors.

### Proposed Model

Previous studies have shown that spontaneous firing rates and somatosensory-evoked neuronal responses decrease in the ZI in a rodent model of central pain syndrome, subsequently disinhibiting the posterior thalamus (PO) (Masri et al., 2009). In the CeA, previous studies have demonstrated that CeA-PKCδ neurons are activated in a model of neuropathic pain and drive behavioral hypersensitivity (Wilson et al., 2019). Given the results presented here demonstrating a strong monosynaptic inhibitory input from the CeA to the ZI (**Figure 3**), in combination with the ZI retrograde tracer experiments showing high colocalization of retrograde tracer uptake in CeA-PKCδ neurons (**Figure 2**) and the behavioral experiments showing that inhibition of ZI GABAergic neurons results in robust bilateral behavioral hypersensitivity (**Figure 4**), we propose a model where injury-induced activation of CeA-PKCδ neurons inhibits ZI-GABAergic cells, subsequently leading to behavioral hypersensitivity (**Figure 8**).

**Figure 8.**
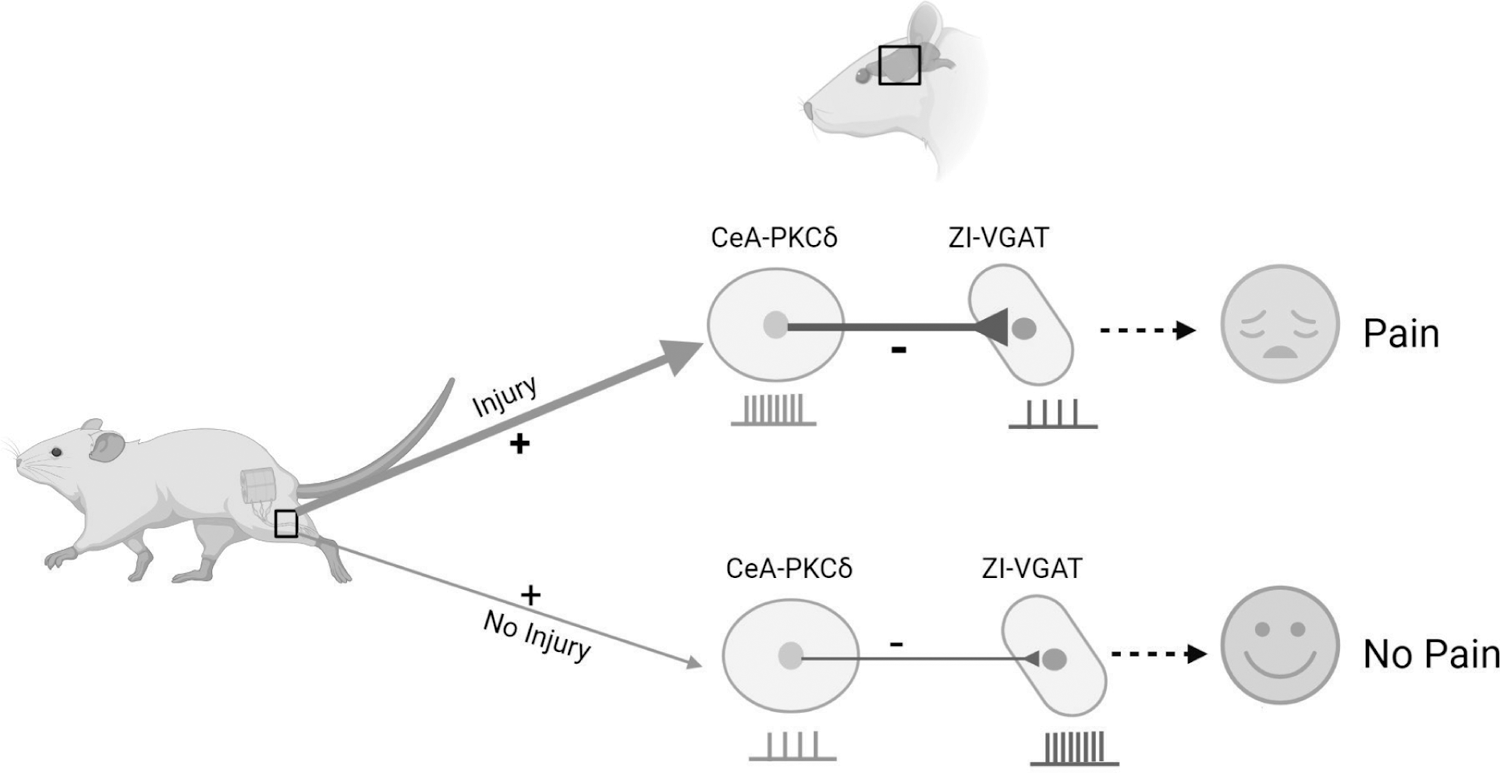
Proposed model for modulation of pain-related behaviors in the ZI. Schematic drawing showing ZI-VGAT neurons receive monosynaptic inhibitory inputs from CeA-PKCδ neurons and modulate pain-related behaviors. After injury, activation of CeA-PKCδ neurons inhibits ZI-VGAT neurons, subsequently leading to behavioral hypersensitivity.

### CeA inputs to Zona Incerta

Our optogenetic-assisted circuit mapping experiments showed that both VGAT-positive and VGAT-negative ZI cells respond to optogenetic stimulation of CeA terminals in the ZI (**Figure 3**). Similarly, our anatomical experiments show retrograde uptake in both PKCδ-positive and PKCδ-negative cells in the CeA when the retrograde tracer was injected in the ZI (**Figure 2**). These results are consistent with previous studies that have shown that somatostatin-expressing neurons in the CeA, which have virtually no overlap with CeA-PKCδ neurons, also project to the ZI (Zhou et al., 2018). Interestingly, in this study, Zhou and colleagues further demonstrated that efferent inputs from somatostatin-expressing neurons in the CeA selectively target parvalbumin-expressing neurons in the ZI and contribute to fear memory. Together, these results suggest that genetically distinct CeA neurons might target neurochemically distinct subpopulations of neurons in the ZI and that cell-type-specific CeA-ZI pathways might differentially contribute to the modulation of pain-related behaviors.

### Modulation of pain-related behaviors in the ZI

In the present study, we show that chemogenetic manipulation of the activity of VGAT-positive ZI neurons bidirectionally modulate pain-related behaviors in a modality-specific manner (**Figures 4-7**). Thus, we observe bilateral hypersensitivity to tactile (but not thermal) stimulation following inhibition of VGAT-positive ZI neurons in the absence of nerve or tissue injury (**Figures 4-5**). In contrast, chemogenetic activation of VGAT-positive ZI neurons reversed nerve injury-induced hypersensitivity to pinch (but not thermal) stimulation. Our results are consistent with previous studies that have shown that optogenetic activation or inhibition of VGAT-positive ZI neurons influences sensitivity to tactile stimulation (Hu et al., 2019). Notably, the behavioral effects of chemogenetic inhibition of the ZI are similar to those observed after chemogenetic activation of the CeA-PKCδ neurons, which also results in increases in tactile (but not thermal) sensitivity (Wilson et al., 2019). Together, these results support our proposed model where injury-induced activation of CeA-PKCδ neurons serves as the source of inhibition of ZI, contributing to behavioral hypersensitivity (**Figure 8**).

Interestingly, the behavioral effects of chemogenetic inhibition of CeA-PKCδ neurons were only partially replicated by the activation of ZI GABAergic neurons. Thus, while inhibition of CeA-PKCδ neurons reversed cuff-induced hypersensitivity to tactile, cold and heat hypersensitivity (Wilson et al., 2019), chemogenetic activation of ZI GABAergic neurons only reversed cuff-induced pinch, but not cold or thermal, hypersensitivity (**Figure 6-7**). Previous studies have shown that responses of spinoparabrachial neurons are modality-specific (Hachisuka et al., 2020). Our results raise the interesting possibility that subpopulations CeA-PKCδ neurons might also be recruited in a modality-specific manner, with CeA-PKCδ neurons that project to the ZI selectively responding to tactile but not thermal stimulation.

### Inconsistencies with previous studies and potential cell-type-specificity in the ZI

Baseline responses in sham animals or the hindpaw contralateral to sciatic nerve treatment were unaltered by chemogenetic activation of VGAT-positive ZI neurons in the present study. This contrasts with previous reports, where optogenetic activation of VGAT-positive ZI neurons decreased baseline responses to tactile stimulation (Hu et al., 2019). Previous studies have demonstrated that modulation of behavioral output by optogenetic stimulation is dependent on the frequency and pattern of the stimulation used for the experiments (Padilla-Coreano et al., 2019). It is therefore possible that the differences in the results from our chemogenetic study and the previous optogenetic study might stem from different levels of neuronal activation by these various techniques.

In the present study, we evaluated the functional contribution of VGAT-positive ZI neurons to the modulation of pain-related behaviors. A separate recent study evaluated the contribution of a different subpopulation of ZI neurons-those expressing parvalbumin (Wang et al., 2020). Interestingly, this study demonstrated that parvalbumin-positive neurons in the ZI show an opposite electrophysiological phenotype and function in the modulation of pain-related behaviors than what we and others observe for the VGAT-positive ZI neurons. Thus, while the present and previous studies show that inhibition of VGAT-positive neurons drives behavioral hypersensitivity, activation of parvalbumin-positive ZI neurons have been previously shown to also promote behavioral hypersensitivity (Wang et al., 2020). These results are particularly interesting given that other studies have demonstrated that parvalbumin-positive ZI neurons receive inputs from somatostatin-expressing CeA neurons (Zhou et al., 2018), which have been previously shown to be inhibited in the context of pain (Wilson et al., 2019).

Based on these combined findings, we propose a model where injury-induced inhibition of somatostatin-expressing neurons in the CeA disinhibits parvalbumin-positive ZI neurons, while injury-induced activation of CeA-PKCδ neurons inhibits VGAT-positive ZI neurons, both resulting in behavioral hypersensitivity. Whether injury-induced inhibition of VGAT-positive ZI neurons (by CeA-PKCδ neurons) occurs simultaneously and under the same conditions as disinhibition of parvalbumin-positive ZI neurons (by somatostatin-expressing neurons in the CeA) remains to be determined. Future studies to address these important questions directly will provide further insights about the CeA-ZI neuronal connection and the potential cell-type-specific contributions to the modulation of pain-related behaviors.

## Material and Methods

### Animals

All experiments were approved by the Animal Care and Use Committee of the National Institute of Neurological Disorders and Stroke and the National Institute on Deafness and other Communication Disorders with the guidelines set by the National Institutes of Health (NIH). Adult male mice (8 to 17–weeks old) were used for all experiments. Mice were housed in a vivarium with controlled humidity and temperature under reversed 12 h light/dark cycle (9 pm to 9 am light) with *ad libitum* access to food and water. All the behavioral tests were performed during the dark period, between the hours of 10 am and 6 pm. Mice received 100 µl of saline intraperitoneal injection (i.p.) daily and were handled by the experimenter for one week before the start of behavioral and electrophysiological experiments following the cupping method as previously described (Hurst and West, 2010). Following surgeries, mice were housed in pairs and separated by a perforated Plexiglas divider. Heterozygous male or female *Prkcd*-cre mice (GENSAT-founder line 011559-UCD) were crossed with Ai9 mice (Jackson Laboratories); vesicular GABA transporter Cre mice (*Slc32a1*-ires-Cre, or VGAT-Cre mice; Jackson Laboratories) were bred as homozygous pairs or crossed with Ai9 mice (Jackson Laboratories). Both the *Prkcd*-cre and VGAT-Cre mouse lines have been previously validated and shown to express Cre selectively in PKCδ+ and GABAergic neurons, respectively (Vong et al., 2011; Wilson et. al., 2019). The presence of cre-recombinase in offspring was confirmed by genotyping using DNA extracted from tail biopsies. The primer sequences (Transnetyx) used for genotyping were TTAATCCATATTGGCAGAACGAAAACG (forward primer) and AGGCTAAGTGCCTTCTCTACA (reverse primer).

### Stereotaxic Injections in the CeA and ZI

Mice were initially anesthetized with 5% isoflurane in preparation for the stereotaxic surgery. After induction, mice were head-fixed on a stereotaxic frame (David Kopf Instruments) and 1.5-2% isoflurane at a flow rate of 0.5 L/min was used for the duration of surgery. A hand warmer was used for thermal maintenance during the procedure. Stereotaxic injections were performed using a 32-gauge needle that fits on a 0.5 µl volume Hamilton Neuros syringe. All the injections were performed at a flow rate of 0.1 µl/min and the syringe was left in place for an additional 5 min after injections to allow for the diffusion of virus and to prevent backflow.

For anterograde labeling, 0.3 µl of rAAV8-hSyn-DIO-mCherry (Addgene viral prep #50459-AAV8) or AAV9-Syn-Flex-ChrimsonR-TdTomato (UNC GTC Vector Core, AV4384G) was microinjected into the right CeA of *Prkcd*-cre mice. The coordinates relative to bregma were as follows: AP −1.4 mm, ML + 3.2, DV −4.8 mm. For retrograde labeling, 0.2 µl of Alexa Fluor 647-conjugated cholera toxin subunit B (Invitrogen, C34778) was stereotaxically injected into the right ZI of *Prkcd*-cre::Ai9 mice. The coordinates relative to bregma were as follows: AP −1.9 mm, ML −1.4 mm, DV −4.75 mm. For optogenetic-assisted circuit mapping experiments, 0.3 µl of rAAV2-hSyn-hChR2(H134R)-EYFP-WPRE-PA (UNC GTC Vector Core, AV6556C) was microinjected into the right CeA of VGAT-Cre::Ai9 mice. The coordinates relative to bregma were as follows: AP −1.25 mm, ML −3.0 mm, DV −4.5 mm. For chemogenetic experiments, 0.15 µl of pAAV8-hSyn-DIO-hM4D(Gi)-mCherry (Addgene viral prep #44362-AAV8; Krashes et al., 2011), pAAV8-hSyn-DIO-hM3D(Gq)-mCherry (Addgene viral prep #44361; Krashes et al., 2011) or rAAV8-hSyn-DIO-mCherry (Addgene viral prep #50459-AAV8) were microinjected into the right ZI of VGAT-Cre mice. The coordinates relative to bregma were as follows: AP −1.7 mm, ML −1.2 mm, DV −4.7 mm. A minimum of 2 weeks was given between brain injections and behavior testing to allow for efficient viral-mediated transduction. For retrograde, anterograde and optogenetic experiments, we waited a minimum of 4 weeks between the brain injections and the experiments.

### Sciatic cuff implantation

Sciatic cuff implantation surgeries were performed 1 week after the brain surgeries as previously described (Benbouzid et al., 2008). Briefly, mice were anesthetized with 2% isoflurane at a flow rate of 0.5 L/min. An incision of about 1-cm long was made in the proximal one third of the lateral left thigh. The sciatic nerve was externalized and stretched using forceps. For the cuff-implanted group, a polyethylene tubing PE20 (2 mm-long, 0.38 mm ID / 1.09 mm OD; Daigger Scientific) was slid onto the sciatic nerve and was then placed back in its location. Similarly, for the comparative sham group, mice went through the same process of sciatic nerve exposure and stretching but no tubing was implanted. After the procedure was complete, the skin above the thigh was closed with wound clips. The mice were subjected to at least one-week recovery period before performing the behavior tests.

### Nociceptive Behaviors

Behavioral experiments were done under red light, during the dark phase, and the experimenter was blind to treatment conditions. Mice were randomized into control and experimental groups. The timeline for the behavioral experiments relative to AAV brain injections were as follows: Acetone test: 14-15 days; Hargreaves test: 16-17 days; Von-Frey test: 20-22 days; Randall-Sellito test: 23-25 days. Each test was performed on 2 consecutive days. On each testing day, baseline (pre-injection) measurements were taken. Saline or Clozapine-N-oxide (CNO, Enzo Life Sciences, Farmingdale, NY) was injected i.p. (10 mg/kg body weight for hM4Di and 5 mg/kg for hM3Dq experiments) and a second measurement (post-injection) was taken 45 min to 1 hour after the i.p. injection. Mice were randomly assigned to saline or CNO on the first day of each test and were tested on the opposite treatment on the second day of the same test.

### Acetone Test

Cold hypersensitivity was assessed by the acetone evaporation test as previously described (Choi Yoon 1994). Briefly, mice were habituated for at least 2 hours in individual ventilated opaque white Plexiglas testing chambers (11 x 11 x 13 cm) on an elevated platform with a floor made of wire mesh. An acetone drop was formed at the top of a 1 mL syringe and gently touched to the center of the plantar surface of the hind paw ipsilateral or contralateral to sciatic nerve surgery. Nociceptive responses to the acetone drop were evaluated for 60 seconds using a modified 0-2-point system developed by Colburn et al., 2007. According to this scoring system, 0 = rapid, transient lifting, licking, or shaking of the hind paw, which subsides immediately; 1 = lifting, licking, and/or shaking of the hind paw, which continues beyond the initial application, but subsides within 5 s; 2 = protracted, repeated lifting, licking, and/or shaking of the hind paw. Five trials were performed with ∼5 min between-trial intervals. Data is represented as an average score of 5 stimulations from each hindpaw before, 1 hour and 1 day after drug treatment (CNO or vehicle).

### Hargreaves test

To evaluate heat hypersensitivity, we used a modified version of the Hargreaves Method (Hargreaves et. al., 1988) as previously described (Wilson et.al., 2019). On the experiment day, mice were habituated prior to testing for at least 1 hour in individual ventilated opaque white plexiglass testing chambers (11 x 11 x 13 cm) placed on an elevated glass floor maintained at 30°C. Following habituation, a noxious radiant heat beam was applied through the glass floor (IITC Life Sciences, Woodland Hills, CA) to the center of the plantar surface of the hindpaw (ipsilateral or contralateral to sciatic nerve surgery), until the mouse showed a withdrawal response. A cutoff of 15 seconds latency and 25 active intensity was used in each trial to prevent skin lesions. At least 3 minutes were allowed between consecutive trials. The average of five trials was calculated and used as the threshold for each hindpaw.

### Von Frey

Mechanical hypersensitivity was assessed as the paw withdrawal threshold in response to von Frey filaments (North Coast Medical, Inc. San Jose, CA), as previously described (Wilson et. al., 2019). On each testing day, mice were placed individually in ventilated opaque white Plexiglas testing chambers (11 x 11 x 13 cm) on an elevated mesh platform at least 2 hours before application of stimulus. Mesh floor allowed full access to the paws from below. After the acclimation period, each von Frey filament was applied to the center of the plantar surface of the hindpaw (ipsilateral or contralateral to sciatic nerve surgery) for 2-3 seconds, with enough force to cause slight bending against the paw. This procedure continued for a total of five measurements. The smallest filament that evoked a paw withdrawal response in at least three of five measurements was taken as the mechanical threshold for that trial. The average of five trials was calculated and used as the threshold value per hindpaw.

### Randall Selitto Test

The Randall Selitto test was performed to assess the response thresholds to mechanical pressure stimulation (pinch) of the hind paws (Randall and Selitto, 1957) in lightly anesthetized animals. Briefly, mice were anesthetized in 5% isoflurane in an induction chamber. Subsequently, animals were kept under light anesthesia with 0.5%– 1% isoflurane at a flow rate of 0.5 L/min. A pinch stimulation of less than 200 g of force was delivered to the plantar surface of the hindpaw ipsilateral or contralateral to sciatic nerve treatment. Pinch pressure stimulation was applied at approximately 1-minute intervals for 30 minutes. The pressure that triggered withdrawal of paw was recorded in all trials. The data is represented as the average response for all trials for each animal before, 1 hour and 1 day after drug treatment (CNO or vehicle).

### Slice Electrophysiology

Acute coronal ZI slices were prepared from brains of VGAT-Cre or VGAT-Cre::Ai9 mice (12-to 18-week-old) 2-3 weeks after stereotaxic injection of AAV8-hSyn-DIO-hM4Di-mCherry into the ZI or 5-8 weeks after AAV2-hsyn-hChR2(H134R)-EYFP injection into the CeA. Briefly, mice were deeply anesthetized with 1.25% Avertin anesthesia (2,2,2-tribromoethanol and tert-amyl alcohol in 0.9% NaCl; 0.025 ml/g body weight), sacrificed by cervical dislocation and decapitated. The brains were rapidly removed and placed in an ice-cold cutting solution containing (in mM): 110.0 choline chloride, 25.0 NaHCO_3_,1.25 NaH_2_PO_4_, 2.5 KCl, 0.5 CaCl_2_, 7.2 MgCl_2_, 25 D-glucose,12.7 L-ascorbic acid, 3.1 pyruvic acid, and saturated with 95% O_2_-5% CO_2_. Coronal slices (300 µm) containing the ZI were cut on a Leica VT1200 S vibrating blade microtome (Leica Microsystems Inc., Buffalo Grove, IL, USA) and incubated in a holding chamber with oxygenated artificial cerebral spinal fluid (ACSF) containing (in mM): 125 NaCl, 2.5 KCl, 1.25 NaH_2_PO_4_, 25 NaHCO_3_, 2.0 CaCl_2_,1.0 MgCl_2_, 25 D-glucose (∼310 mOsm) saturated with 95% O_2_-5% CO_2_, at 33° C for 30 minutes, then moved to room temperature for at least an additional 20 minutes before transfer to the recording chamber.

To validate the effects of CNO on hM4Di-transduced cells in the ZI, current-clamp recordings using potassium methylsulfate-based internal solution (in mM: 120 KMeSO_4_, 20 KCl, 10 HEPES, 0.2 EGTA, 8 NaCl, 4 Mg-ATP, 0.3 Tris-GTP, 14 Phosphocreatine, pH 7.3 with KOH (∼300 mosmol-1) were used to assess changes in excitability. Whole-cell current-clamp recordings were obtained at 33 ± 1° C from ZI cells expressing mCherry. A recording chamber heater and an in-line solution heater (Warner Instruments) were used to control and monitor the bath temperature throughout the experiment. Cells were visually identified using an upright microscope (Nikon Eclipse FN1) equipped with differential interference contrast optics with infrared illumination and fluorescent microscopy. Spontaneously active cells were injected with hyperpolarizing current (−10 to −50 pA) to bring their membrane potentials to between −80 and −70 mV. A 500 ms depolarizing current (10 to 120 pA) was injected to elicit between 2 and 5 action potentials. The current injection repeated every 15 seconds until the cell fired stably and consistently. Following this stabilization, 10 additional recordings were acquired before bath application of either 10 µM CNO or vehicle in ACSF. Recordings of the same current injection were continued every 15 seconds for approximately 5 minutes, until the cell fired consistently and stably. Ten additional recordings were performed. The number of action potentials elicited during each depolarizing current injection were used to assess excitability. Values for before and after CNO or vehicle application were averaged across five traces and compared. Current clamp signals were acquired at 100 kHz and filtered at 10 kHz.

### Optogenetically assisted Circuit Mapping

For the optogenetically assisted circuit mapping experiments, recordings were done using a cesium gluconate-based internal solution containing (in mM): 120 cesium gluconate, 6 NaCl, 10 HEPES, 12 phosphocreatine, 5 EGTA, 1 CaCl_2_, 2 MgCl_2_, 2 ATP, and 0.5 GTP, pH 7.4 adjusted with CsOH (∼290 mOsm). Whole-cell voltage-clamp recordings were obtained at 33 ± 1° C from visually identified tdTomato-expressing and non-expressing ZI neurons using differential interference contrast optics with infrared illumination and fluorescence microscopy. Optically evoked inhibitory postsynaptic currents (oIPSCs) were recorded at a holding potential of 0 mV in the presence of TTX (1 µM) and 4-AP (100 µM). Blue LED light (λ = 470 nm, Mightex) paired pulses of 5 ms duration with an interval of 200 ms between pulses were delivered at 10 Hz to drive paired synaptic responses. Paired pulse ratios (PPR) were determined by the ratio of the amplitude of the peak evoked by the second pulse divided by the amplitude of the peak evoked by the first pulse. Signals were acquired at 100 kHz and filtered at 10kHz.

### Immunohistochemistry

At the end of each experiment, mice were deeply anesthetized with 1.25% Avertin anesthesia (2,2,2-tribromoethanol and tert-amyl alcohol in 0.9% NaCl; 0.025 ml/g body weight) i.p., then perfused transcardially with 0.9% NaCl (37°C), followed by 100 mL of ice-cold 4% paraformaldehyde in phosphate buffer solution (PFA/PB). The brain was dissected and post fixed in 4% PFA/PB overnight at 4°C. After cryoprotection in 30% sucrose/PB for 48 h, coronal sections (30-45 μm) were obtained using a freezing sliding microtome and stored in 0.1 M Phosphate Buffered Saline (PBS), pH 7.4 containing 0.01% sodium azide (Sigma) at 4°C until immunostaining. Sections were rinsed in PBS, incubated in 0.1% Triton X-100 in PBS for 10 minutes at room temperature and blocked in 5% normal goat serum (NGS) (Vector Labs, Burlingame, CA) with 0.1% Triton X-100, 0.05% Tween-20 and 1% bovine serum albumin (BSA) for 30 minutes at room temperature. Sections were then incubated for 72 h at 4°C in mouse anti-PKCδ (1:1000, BD Biosciences, 610397), rabbit anti-Phospho-c-Fos (1:2000, Cell Signaling Technology, 5348) or rat anti-mCherry (1:500, Invitrogen, M11217) in 1.5% NGS blocking solution with 0.1% Triton X-100, 0.05% Tween-20 and 1% BSA. Sections were then rinsed in PBS and incubated in Alexa Fluor 647-conjugated goat anti-mouse (1:100, Invitrogen, A21235), Alexa Fluor 647-conjugated goat anti-rabbit (1:250, Invitrogen, A21244), or goat anti rat Cy3 (1:250, Invitrogen, A10522) secondary antibodies in 1.5% NGS blocking solution with 0.1% Triton X-100, 0.05% Tween 20 and 1% BSA, protected from light, for 2 h at room temperature. Sections were then rinsed in PBS, mounted on positively charged glass slides, air-dried and coverslips were placed using Fluoromount-G (Southern Biotech).

For the c-Fos experiments, VGAT-Cre mice received CNO (5 mg/kg) or saline injections (i.p) 2 weeks post virus injection into the ZI. Mice were housed in their home cages for 1 hour prior to transcardial perfusion, brain dissection and tissue processing. For the mapping of axonal terminals from CeA-PKCδ neurons, PKCδ-cre mice injected with AAV9-Syn-Flex-ChrimsonR-TdTomato or AAV8-hSyn-DIO-mCherry were transcardially perfused at least 4 weeks after the brain injections. For the mCherry-injected brains, 30 µm coronal sections from the entire brain where collected and immunostained for mCherry as described above. No staining was performed for the Chrimson-R-injected brains.

### Imaging and analysis

For confocal studies, images were acquired using a Nikon A1R laser scanning confocal microscope. 2X (for low magnification), 20X (for high magnification) or 40X (oil-immersion for higher magnification) objectives were used. The experimental conditions for image collection including laser intensity, gain, and pinhole were identical for experiments. Multiple channels (GFP, RFP and CY5) were used for sequential image acquisition where Z stacks data collection was done at 0.9 mm. Following acquisition, images were consolidated using NIS Elements software with automatic stitching of subsequent images, and conversion of stacks into maximum intensity z-projections. Quantitative analysis of CeA and ZI imaging data was performed between bregma −0.82 and −1.82 and bregma −1.06 and −2.54, for CeA and ZI, respectively. Anatomical limits of each region were identified based on the mouse brain atlas (Paxinos and Franklin, 2001). Number of positive cells were quantified manually for each channel using NIS Elements software using one section per rostro caudal level for each mouse. Co-labeled cells were identified by NIS Elements software automatically and were further visually corroborated by an experimenter.

For the mapping of axonal terminals from CeA-PKCδ neurons, coronal slices from the entire brain of PKCδ-cre mice injected with AAV9-Syn-Flex-ChrimsonR-TdTomato or AAV8-hSyn-DIO-mCherry, collected and immunostained 120 µm apart from each other, were visually inspected for the presence of fluorescent axonal terminals using a 20X objective in a Nikon A1R laser scanning confocal microscope. Classic morphological criteria, defined as the presence of varicosities and the thickness and organization pattern of the signal, was used to distinguish labeled terminals (very thin fibers with numerous ramifications and varicosities) from fibers of passage (thicker fibers without ramifications and varicosities) as previously described (Bernard et al.,1993). Low and high magnification images of all the brain sections containing terminals were acquired and the anatomical localization of the terminals was then determined using a Mouse Brain Atlas (Paxinos and Franklin, 2001) Representative images of terminals were collected using a 40X oil-immersion objective.

Semi-quantitative analysis of the areas containing axonal terminals was done based on the density of terminals observed and are reported as sparce (+), moderate (++) and dense (+++) in the defined area. A similar analysis was also performed with the brain of experiment 265945645 of the Mouse Brain Connectivity Atlas of the Allen Brain Institute (http://connectivity.brain-map.org/). This brain was identified using the Source Search tool on the Mouse Brain Connectivity Atlas website and filtering for the CeA as the brain region and *Prkcd*-GluCla-CFP-IRES-Cre as the mouse line of interest. The tracer injected is described as EGFP; the stereotaxic coordinates for the injection were AP −1.82 mm, ML −2.65 mm, DV −4.25 mm and the injection volume was 0.02 mm^3^.

### Statistical Analysis

Data are presented as mean ± SEM. Statistical analyses was conducted using GraphPad Prism (v8.0). Unpaired/paired two-tailed t-test, Wilcoxon two-tailed matched pair signed rank test, one-way analyses of variance (ANOVA) followed by Tukey’s multiple comparison test and two-way ANOVA followed by Dunnett’s multiple comparison test were used. The significance level was set at p < 0.5. Sample sizes and p values are indicated in figure legends. Detailed information on each statistical test performed are shown in **Supplemental Table 1**.

## Acknowledgements

This research was supported by the National Center for Complementary and Integrative Health Intramural Research Program. The authors would like to thank Simón Arango, Adela Francis Malavé and Jeitzel Torres Rodriguez for their assistance in histological and anatomical experiments. pAAV-hSyn-DIO-hM3D(Gq)-mCherry (Addgene viral prep # 44361-AAV8), pAAV-hSyn-DIO-hM4D(Gi)-mCherry (Addgene viral prep # 44362-AAV8), and pAAV-hSyn-DIO-mCherry (Addgene viral prep # 50459-AAV8) were a gift from Bryan Roth. The AAV2-hsyn-hChR2(H134R)-EYFP was provided by the Vector Core at the University of North Carolina, with a material transfer agreement with Professor Karl Deisseroth (Stanford University).

---

**Figure 1 – figure supplement 1.**
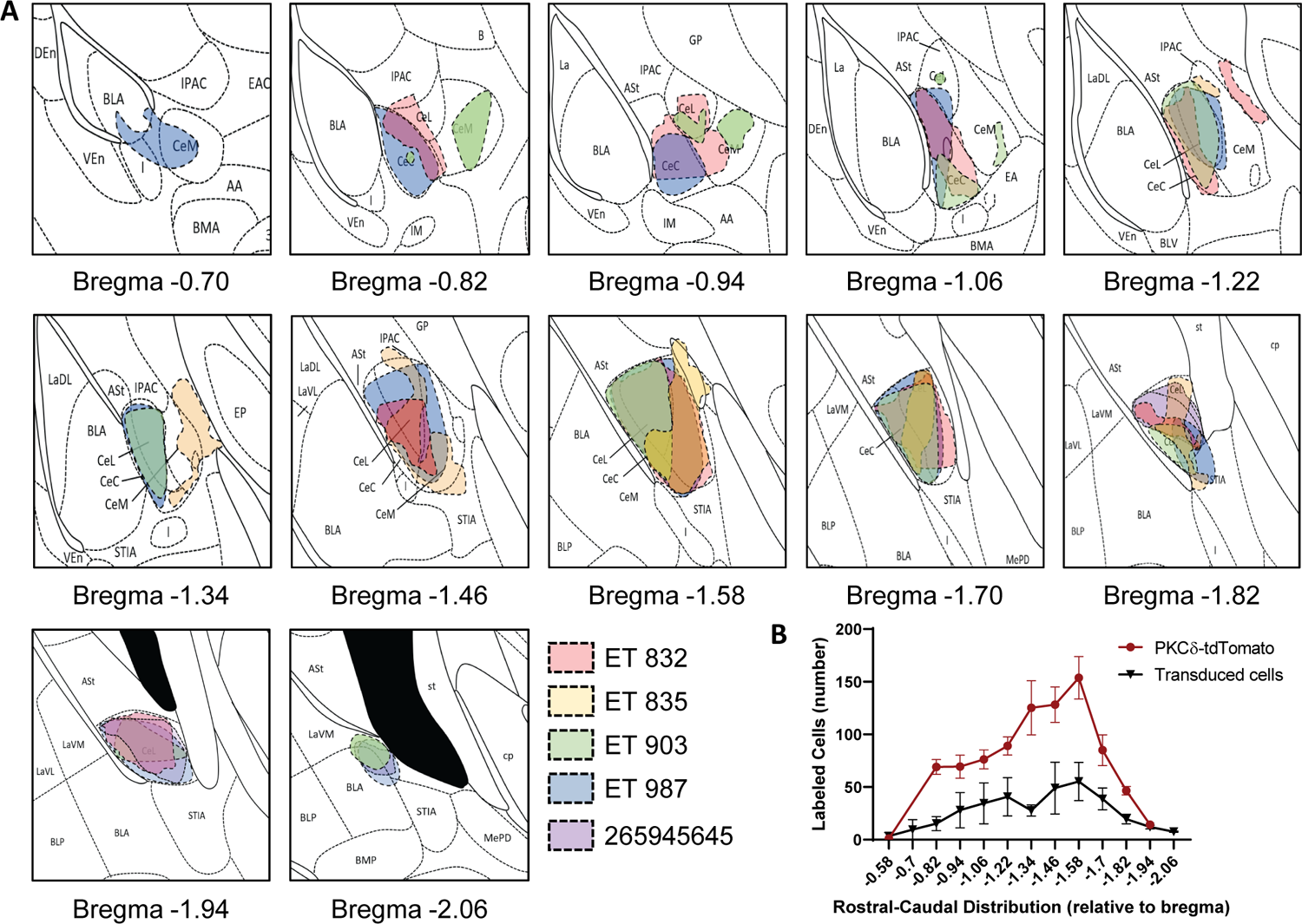
(**A**) Rostro-caudal distribution of AAV8-hSyn-DIO-mCherry or AAV1-Syn-Flex-ChrimsonR-tdTomato injection sites in mice used for anatomical analysis. Drawings of injection sites throughout *Prkcd*-Cre mice brains injected into the CeA with AAV8-hSyn-DIO-mCherry (ET 832 and ET835), AAV1-Syn-Flex-ChrimsonR-tdTomato (ET 903 and ET 987) or EGFP (experiment 265945645 of the Mouse Brain Connectivity Atlas of the Allen Brain Institute-http://connectivity.brain-map.org/). Individual mice are represented in different colors. (**B**) Mean ± SEM number of neurons transduced with Chrimson-R-tdTomato or mCherry and PKCδ-tdTomato labeled cells in the CeA as a function of rostro-caudal level relative to bregma (n=4 mice for transduced neurons; n=5 mice for PKCδ-tdTomato neurons).

**Figure 4 – figure supplement 1.**
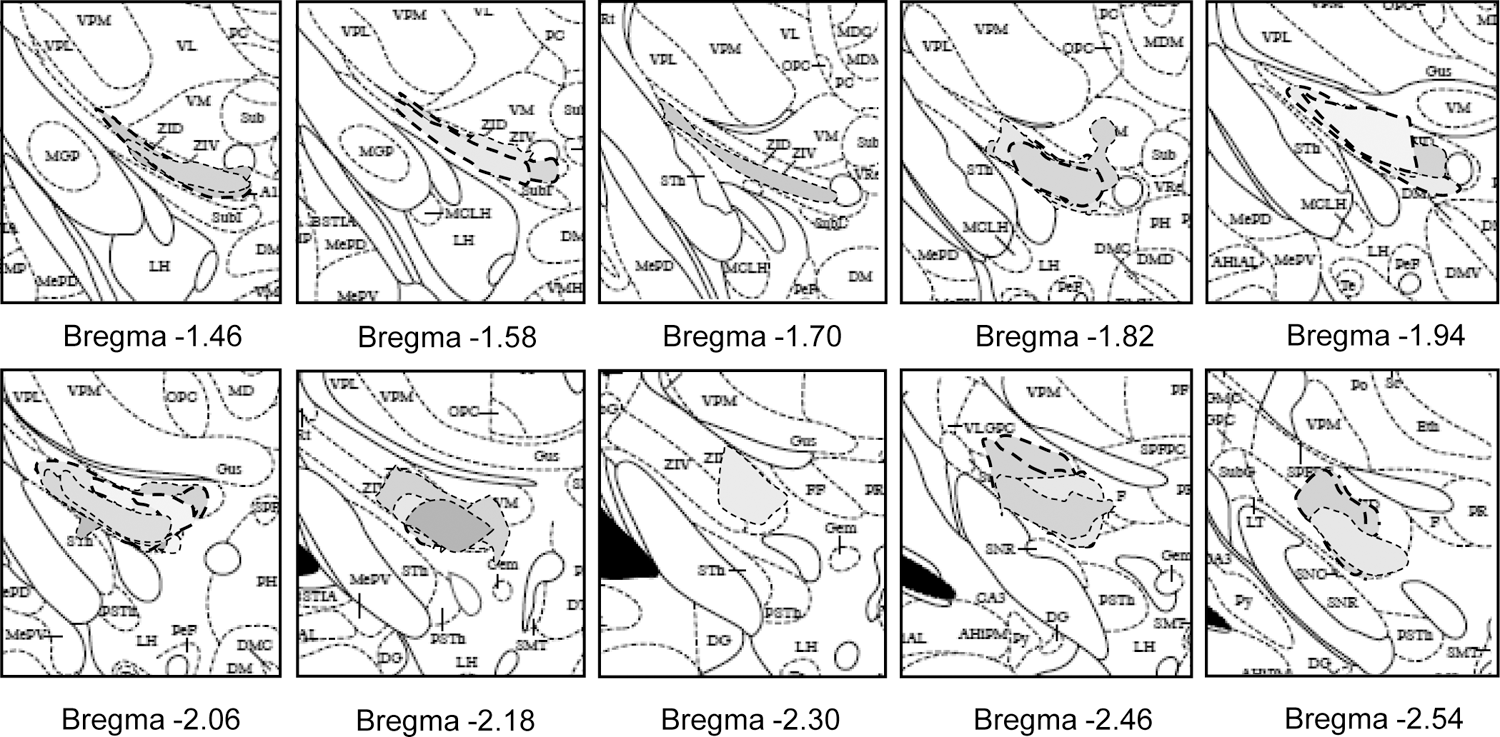
Rostro-caudal distribution AAV-hSyn-DIO-hM4D(Gi)-mCherry injection sites in mice used for behavioral experiments. Drawings of injection sites throughout the VGAT-Cre mice brains injected with AAV-hSyn-DIO-hM4D(Gi)-mCherry into the ZI. Individual mice are represented in different color.

**Figure 6 – supplemental figure 1.**
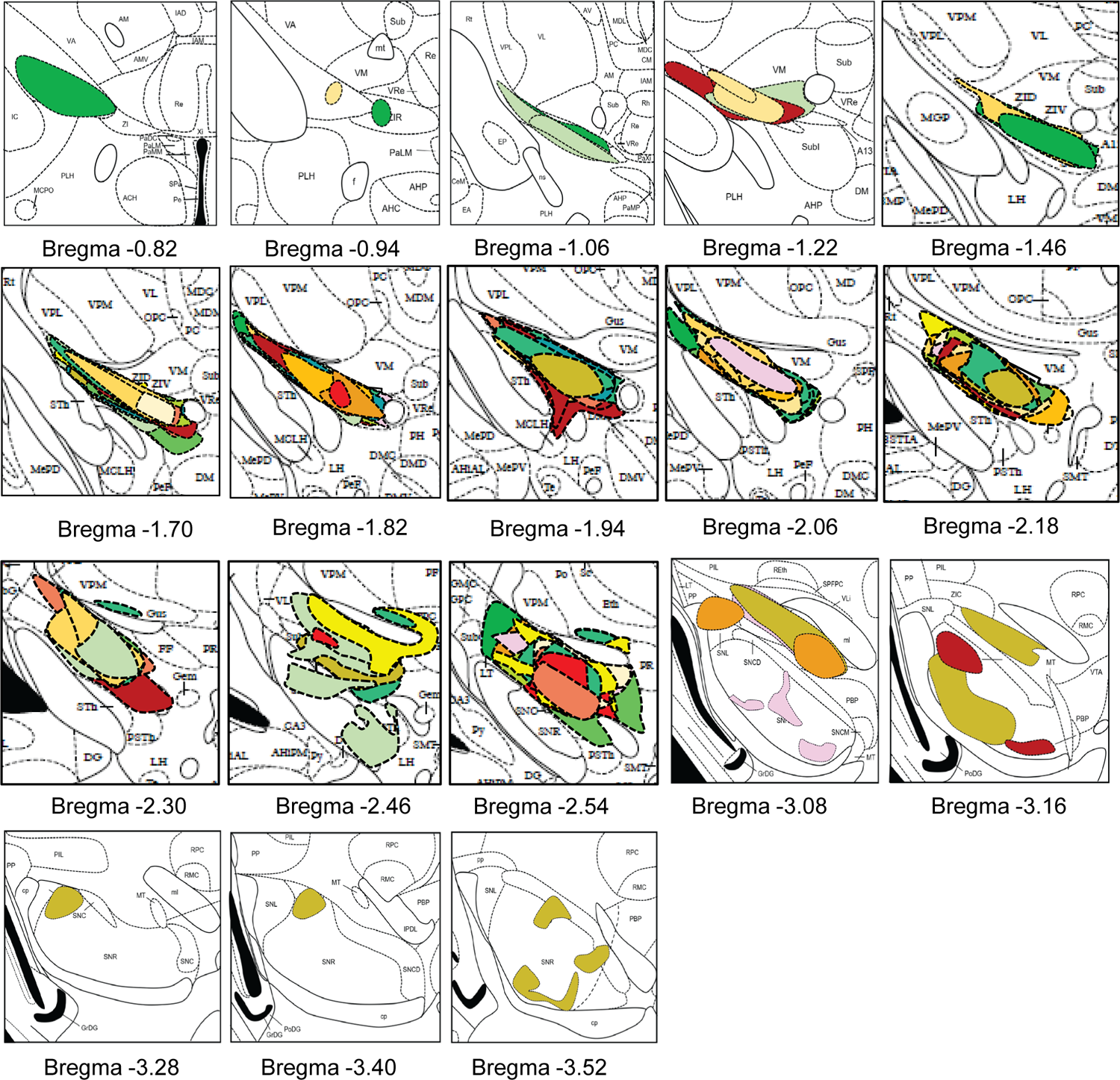
Rostro-caudal distribution of AAV-hSyn-DIO-hM3D(Gq)-mCherry injection sites in mice used for behavioral experiments. Drawings of injection sites throughout the VGAT-Cre mice brains injected with AAV-hSyn-DIO-hM3D(Gq)-mCherry into the ZI. Individual mice are represented in different color.

**Figure 6 – supplemental figure 2.**
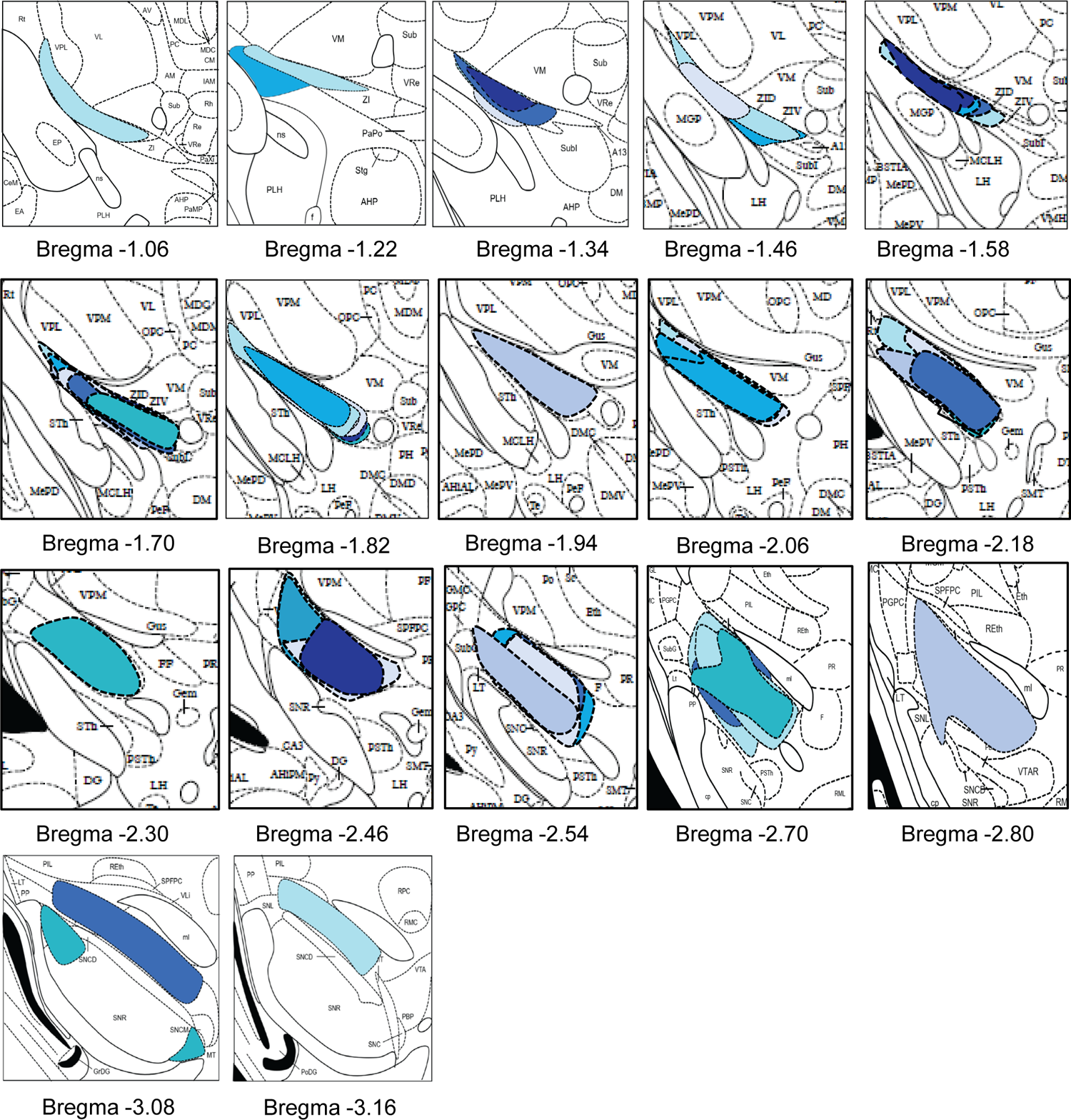
Rostro-caudal distribution of AAV8-hSyn-DIO-mCherry injection sites in mice used for behavioral experiments. Drawings of injection sites throughout the VGAT-Cre mice brains injected with AAV8-hSyn-DIO-mCherry into the ZI. Individual mice are represented in different color.

**Supplemental Table 1.** Statistical Analyses Detailed information about data structure, statistical tests, sample sizes and statistical results; ANOVA = analyses of variance.(DFFn, DFd): degree of freedom for the numerator of the F ratio, for the denominator of the F ratio; df: degrees of freedom.

| Figure | Data Structure | Type of Test | Sample Size | Statistical Data |
| --- | --- | --- | --- | --- |
| **Figure 3** |  |  |  |  |
| 3D (Ex vivo Optogenetics) | Normal distribution | Unpaired two-tailed t-test | VGAT-positive = 8 cells<br>VGAT-negative = 12 cells | t = 0.3291; df = 18; p = 0.7459; eta squared = 0.005982 |
| **Figure 4** |  |  |  |  |
| 4A (Gi validation) | Normal distribution<br>Non-normal distribution | Paired two-tailed t-test<br>Wilcoxon two tailed matched-paired signed rank test | Saline = 6 cells<br>CNO = 6 cells | t = 0.98; df = 5; p = 0.3722; eta squared = 0.161<br>p = 0.0313; W = -21.0 |
| 4E (Von Frey) | Normal distribution | Two-way ANOVA followed by Dunnett's multiple comparison test | n = 8 mice per group | Two-way ANOVA: Ipsilateral Paw sciatic nerve and brain treatments: F(1,35) = 0.3491; p = 0.554<br>sciatic nerve and brain treatments: F(1,35) = 188.9; p < 0.000<br>posthoc: Dunnett's test<br>***p < 0.0001 for pre-injections vs 1 h after CNO in sham-hM4Di-CNO treatment<br>Two-way ANOVA: Contralateral Paw i.p. treatment: F(2,35) = 96.25; p < 0.000<br>sciatic nerve and brain treatments: F(1,35) = 0.3491; p = 0.554<br>posthoc: Dunnett's test<br>***p < 0.0001 and ***p < 0.0001 for pre-injections vs 1 h after CNO in sham-hM4Di-CNO and cuff-hM4Di-CNO treatment respectively |
| 4F (Pinching) | Normal distribution | Two-way ANOVA followed by Dunnett's multiple comparison test | Ipsilateral paw-Sham = 8 mice per group<br>Cuff = 6 mice per group<br>Contralateral paw-Sham = 6 mice per group<br>Cuff = 7 mice per group | Two-way ANOVA: Ipsilateral Paw i.p. treatment: F(2,29) = 59.13; p < 0.0001<br>sciatic nerve and brain treatments: F(1,29) = 338.7; p < 0.000<br>posthoc: Dunnett's test<br>***p < 0.0001 for pre-injections vs 1 h after CNO in sham-hM4Di-CNO treatment<br>Two-way ANOVA: Contralateral Paw i.p. treatment: F(2,28) = 210.0; p < 0.0001<br>sciatic nerve and brain treatments: F(1,28) = .2574; p = 0.615<br>posthoc: Dunnett's test<br>***p < 0.0001 and ***p < 0.0001 for pre-injections vs 1 h after CNO in sham-hM4Di-CNO and cuff-hM4Di-CNO treatment respectively |
| **Figure 5** |  |  |  |  |
| 5A (Acetone) | Normal distribution | Two-way ANOVA followed by Dunnett's multiple comparison test | n = 8 mice per group | Two-way ANOVA: Ipsilateral Paw i.p. treatment: F(2,34) = 0.2743; p = 0.7611<br>sciatic nerve and brain treatments: F(1,34) = 0.0046; p = 0.945<br>posthoc: Dunnett's test<br>p = 0.9871 for pre-injections vs 1 h after CNO in sham-hM4Di-CNO treatment<br>Two-way ANOVA: Contralateral Paw i.p. treatment: F(2,32) = 0.018; p = 0.982<br>sciatic nerve and brain treatments: F(1,32) = 1.071; p = 0.308<br>posthoc: Dunnett's test<br>p > 0.9999 and p = 0.9790 for pre-injections vs 1 h after CNO in sham-hM4Di-CNO and cuff-hM4Di-CNO treatment respectively |
| 5B (Hargreaves) | Normal distribution | Two-way ANOVA followed by Dunnett's multiple comparison test | n = 8 mice per group | Two-way ANOVA: Ipsilateral Paw i.p. treatment: F(2,36) = 0.1659; p = 0.8471<br>sciatic nerve and brain treatments: F(1,36) = 0.8412; p = 0.773<br>posthoc: Dunnett's test<br>p = 0.9089 for pre-injections vs 1 h after CNO in sham-hM4Di-CNO treatment<br>Two-way ANOVA: Contralateral Paw i.p. treatment: F(2,37) = 1.822; p = 0.1761<br>sciatic nerve and brain treatments: F(1,37) = 0.07605; p = 0.784<br>posthoc: Dunnett's test<br>p = 0.8940 and p = 0.9040 for pre-injections vs 1 h after CNO in sham-hM4Di-CNO and cuff-hM4Di-CNO treatment respectively |
| **Figure 6** |  |  |  |  |
| 6E (cFos) | Normal distribution: HM3Dq-CNO and mCherry-CNO<br>Non-normal distribution: HM3Dq-Saline | One-way ANOVA followed by Tukey's multiple comparisons test | HM3Dq-CNO = 7 mice<br>HM3Dq-Saline = 5 mice<br>mCherry-CNO = 3 mice | One-way ANOVA: F(2,12) = 10.90; p < 0.001<br>posthoc: Tukey's test<br>**p < 0.01 for HM3Dq-CNO vs HM3Dq-Sal<br>*p < 0.05 for HM3Dq-CNO vs mCherry-CNC |
| 6F (Pinching) | Normal distribution | Two-way ANOVA followed by Dunnett's multiple comparison test | HM3Dq-Sham = 6 mice per group; HM3Dq-Cuff = 8 mice per group; mCherry-Cuff = 8 mice | Two-way ANOVA: Ipsilateral Paw i.p. treatment: F(2,38) = 47.13; p < 0.000<br>sciatic nerve and brain treatments: F(1,38) = 31.18; p < 0.000<br>posthoc: Dunnett's test<br>***p < 0.0001 for pre-injections vs 1 h after CNO in cuff-HM3Dq-CNO treatment<br>Two-way ANOVA: Contralateral Paw i.p. treatment: F(2,38) = 1.245; p = 0.2991<br>sciatic nerve and brain treatments: F(1,38) = 2.069; p = 0.158<br>posthoc: Dunnett's test<br>p = 0.0719 for pre-injections vs 1 h after CNO in cuff-HM3Dq-CNO treatment |
| **Figure 7** |  |  |  |  |
| 7A (Acetone) | Normal distribution | Two-way ANOVA followed by Dunnett's multiple comparison test | HM3Dq-Sham = 6 mice per group; HM3Dq-Cuff = 8 mice per group; mCherry-Cuff = 8 mice | Two-way ANOVA: Ipsilateral Paw i.p. treatment: F(2,34) = 0.1027; p = 0.902<br>sciatic nerve and brain treatments: F(1,34) = 0.5246; p = 0.473<br>posthoc: Dunnett's test<br>p = 0.7548 for pre-injections vs 1 h after CNO in cuff-HM3Dq-CNO treatment<br>Two-way ANOVA: Contralateral Paw i.p. treatment: F(2,34) = 0.4027; p = 0.671<br>sciatic nerve and brain treatments: F(1,34) = 0.2575; p = 0.873<br>posthoc: Dunnett's test<br>p = 0.9363 for pre-injections vs 1 h after CNO in cuff-HM3Dq-CNO treatment |
| 7B (Hargreaves) | Normal distribution | Two-way ANOVA followed by Dunnett's multiple comparison test | HM3Dq-Sham = 6 mice per group; HM3Dq-Cuff = 8 mice per group; mCherry-Cuff = 8 mice | Two-way ANOVA: Ipsilateral Paw i.p. treatment: F(2,14) = 0.9272; p = 0.418<br>sciatic nerve and brain treatments: F(1,14) = 1.036; p = 0.325<br>posthoc: Dunnett's test<br>p = 0.6771 for pre-injections vs 1 h after CNO in cuff-HM3Dq-CNO treatment<br>Two-way ANOVA: Contralateral Paw i.p. treatment: F(2,14) = 1.057; p = 0.373<br>sciatic nerve and brain treatments: F(1,14) = 1.722; p = 0.210<br>posthoc: Dunnett's test<br>p = 0.5475 for pre-injections vs 1 h after CNO in cuff-HM3Dq-CNO treatment |

## References

1. Dworkin RH. An overview of neuropathic pain: syndromes, symptoms, signs, and several mechanisms. Clin J Pain. 2002;18(6):343–349 10.1097/00002508-200211000-00001.

2. Treede RD, Jensen TS. Campbell, J.N. Cruccu, G. Dostrovsky, J.O. Griffin, J.W. Hansson, P. Hughes, R. Nurmikko, T. Serra, J. Neuropathic pain: redefinition and a grading system for clinical and research purposes. Neurology. 2008;70(18):1630–1635 10.1212/01.wnl.0000282763.29778.59.

3. Bernard JF, Besson, JM. The spino(trigemino)pontoamygdaloid pathway: electrophysiological evidence for an involvement in pain processes. J Neurophysiol. 1990;63(3):473–490 10.1152/jn.1990.63.3.473.

4. Carrasquillo Y, Gereau RW 4th. Activation of the extracellular signal-regulated kinase in the amygdala modulates pain perception. J Neurosci. 2007;27(7):1543–1551 10.1523/JNEUROSCI.3536-06.2007.

5. Bushnell MC, Ceko M, Low LA. Cognitive and emotional control of pain and its disruption in chronic pain. Nat Rev Neurosci. 2013;14(7):502–511 10.1038/nrn3516.

6. Neugebauer V, Li W, Bird GC, Han JS. The amygdala and persistent pain. Neuroscientist. 2004;10(3):221–234 10.1177/1073858403261077.

7. Zald DH. The human amygdala and the emotional evaluation of sensory stimuli. Brain Research Reviews 2003;41:88–123. 10.1016/s0165-0173(02)00248-5.

8. Wilson TD, Valdivia S, Khan A, Ahn HS, Adke AP, Martinez Gonzalez S, Sugimura YK, Carrasquillo Y. Dual and Opposing Functions of the Central Amygdala in the Modulation of Pain. Cell Rep. 2019;29(2):332–346 e335 10.1016/j.celrep.2019.09.011.

9. Ricardo JA. Efferent connections of the subthalamic region in the rat. II. The zona incerta. Brain Research. 1981;214(1):43–60 10.1016/0006-8993(81)90437-6.

10. Mitrofanis J. Some certainty for the “zone of uncertainty”? Exploring the function of the zona incerta. Neuroscience. 2005;130(1):1–15 10.1016/j.neuroscience.2004.08.017.

11. Zhou M, Liu Z, Melin MD, Ng YH, Xu W, Sudhof TC. A central amygdala to zona incerta projection is required for acquisition and remote recall of conditioned fear memory. Nat Neurosci. 2018;21(11):1515–1519 10.1038/s41593-018-0248-4.

12. Zhang X, Van Den Pol A. Rapid binge-like eating and body weight gain driven by zona incerta GABA neuron activation. Science. 2017;356(6340):853–859 10.1126/science.aam7100.

13. Chou XL, Wang X, Zhang ZG, Shen L, Zingg B, Huang J, Zhong W, Mesik L, Zhang LI, Tao HW. Inhibitory gain modulation of defense behaviors by zona incerta. Nat Commun. 2018;9(1):1151 10.1038/s41467-018-03581-6.

14. Zhao ZD, Chen Z, Xiang X, Hu M, Xie H, Jia X, Cai F, Cui Y, Chen Z, Qian L, Liu J, Shang C, Yang Y, Ni X, Sun W, Hu J, Cao P, Li H, Shen WL. Zona incerta GABAergic neurons integrate prey-related sensory signals and induce an appetitive drive to promote hunting. Nat Neurosci. 2019;22(6):921–932 10.1038/s41593-019-0404-5.

15. Masri R, Quiton RL, Lucas JM, Murray PD, Thompson SM, Keller A. Zona incerta: a role in central pain. J Neurophysiol. 2009;102(1):181–191 10.1152/jn.00152.2009.

16. Wang H, Dong P, He C, Feng XY, Huang Y, Yang WW, Gao HJ, Shen XF, Lin S, Cao SX, Lian H, Chen J, Yan M, Li XM. Incerta-thalamic Circuit Controls Nocifensive Behavior via Cannabinoid Type 1 Receptors. Neuron. 2020;107(3):538–551 e537 10.1016/j.neuron.2020.04.027.

17. Petronilho A, Reis GM, Dias QM, Fais RS, Prado WA. Antinociceptive effect of stimulating the zona incerta with glutamate in rats. Pharmacol Biochem Behav. 2012;101(3):360–368 10.1016/j.pbb.2012.01.022.

18. Moon HC, Park YS. Reduced GABAergic neuronal activity in zona incerta causes neuropathic pain in a rat sciatic nerve chronic constriction injury model. J Pain Res. 2017;10:1125–1134 10.2147/JPR.S131104.

19. Hu TT, Wang RR, Du Y, Guo F, Wu YX, Wang Y, Wang S, Li XY, Zhang SH, Chen Z. Activation of the Intrinsic Pain Inhibitory Circuit from the Midcingulate Cg2 to Zona Incerta Alleviates Neuropathic Pain. J Neurosci. 2019;39(46):9130–9144 10.1523/JNEUROSCI.1683-19.2019.

20. Moon HC, Lee YJ, Cho CB, Park YS. Suppressed GABAergic signaling in the zona incerta causes neuropathic pain in a thoracic hemisection spinal cord injury rat model. Neurosci Lett. 2016;632:55–61 10.1016/j.neulet.2016.08.035.

21. Hurst JL, West RS. Taming anxiety in laboratory mice. Nat Methods. 2010;7(10):825–826 10.1038/nmeth.1500.

22. Vong L, Ye C, Yang Z, Choi B, Chua S, Jr., Lowell BB. Leptin action on GABAergic neurons prevents obesity and reduces inhibitory tone to POMC neurons. Neuron. 2011;71(1):142–154 10.1016/j.neuron.2011.05.028.

23. Krashes MJ, Koda S, Ye C, Rogan SC, Adams AC, Cusher DS, Maratos-Flier E, Roth BL, Lowell BB. Rapid, reversible activation of AgRP neurons drives feeding behavior in mice. J Clin Invest. 2011;121(4):1424–1428 10.1172/JCI46229.

24. Benbouzid M, Pallage V, Rajalu M, Waltisperger E, Doridot S, Poisbeau P, Freund-Mercier MJ, Barrot M. Sciatic nerve cuffing in mice: a model of sustained neuropathic pain. Eur J Pain. 2008;12(5):591–599 10.1016/j.ejpain.2007.10.002.

25. Choi Y, Wook Y, Yoon Na Heung Sik, Kim Sun Ho, Chung Jin Mo. Behavioral signs of ongoing pain and cold allodynia in a rat model of neuropathic pain. Pain. 1994;59(3):369–376 10.1016/0304-3959(94)90023-X.

26. Colburn RW, Lubin ML, Stone DJ, Jr, Wang Y, Lawrence D, D’Andrea MR, Brandt MR, Liu Y, Flores CM, Qin N. Attenuated cold sensitivity in TRPM8 null mice. Neuron. 2007;54:379–386. 10.1016/j.neuron.2007.04.017.

27. Hargreaves K, Dubner R, Brown F, Flores C, Joris J. A new and sensitive method for measuring thermal nociception in cutaneous hyperalgesia. Pain. 1988;32:77–88 10.1016/0304-3959(88)90026-7.

28. RANDALL LO, SELITTO JJ. A method for measurement of analgesic activity on inflamed tissue. Arch Int Pharmacodyn Ther. 1957;111(4):409–419,

29. Paxinos G, Franklin K.B.J. The Mouse Brain in Stereotaxic Coordinates.: Academic Press; 2001.

30. Bernard JF, Alden M., Besson, J.M. The Organization of the Efferent Projections From the Pontine Parabrachial Area to the Amygdaloid Complex: A Phaseolus vulgaris Leucoagglutinin (PHA-L) Study in the Rat. J Comp Neurol. 1993;329(2):201–229 10.1002/cne.903290205.

31. Reardon F, Mitrofanis J. Organisation of the amygdalo-thalamic pathways in rats. Anat Embryol (Berl). 2000;201:75–84 10.1007/pl00008229.

32. Barbier M, Chometton S, Peterschmitt Y, Fellmann D, Risold PY. Parasubthalamic and calbindin nuclei in the posterior lateral hypothalamus are the major hypothalamic targets for projections from the central and anterior basomedial nuclei of the amygdala. Brain Struct Funct. 2017;222(7):2961–2991 10.1007/s00429-017-1379-1.

33. Shinonaga YT, M. Mizuno, N. Direct projections from the central amygdaloid nucleus to the globus pallidus and substantia nigra in the cat. Neuroscience. 1992;51(3) 10.1016/0306-4522(92)90308-O.

34. Aggleton PJ. The Amygdala A Functional Analysis Second Edition: Oxford University Press; 2nd edition; 2000 21 December 2000. 712 p,

35. Hachisuka J, Koerber HR, Ross SE. Selective-cold output through a distinct subset of lamina I spinoparabrachial neurons. Pain. 2020;161(1):185–194 10.1097/j.pain.0000000000001710.

36. Padilla-Coreano N, Canetta S, Mikofsky RM, Alway E, Passecker J, Myroshnychenko MV, Garcia-Garcia AL, Warren R, Teboul E, Blackman DR, Morton MP, Hupalo S, Tye KM, Kellendonk C, Kupferschmidt DA, Gordon JA. Hippocampal-Prefrontal Theta Transmission Regulates Avoidance Behavior. Neuron. 2019;104(3):601–610 e604 10.1016/j.neuron.2019.08.006.

